# Single-cell time series analysis reveals the dynamics of *in vivo* HSPC responses to inflammation

**DOI:** 10.1101/2023.03.09.531881

**Authors:** Brigitte Joanne Bouman, Yasmin Demerdash, Shubhankar Sood, Florian Grünschläger, Franziska Pilz, Abdul Rahman Itani, Andrea Kuck, Simon Haas, Laleh Haghverdi, Marieke Alida Gertruda Essers

**Author notes:** These authors contributed equally to this work (Brigitte Joanne Bouman, Yasmin Demerdash).

## Abstract

Hematopoietic stem and progenitor cells (HSPCs) are known to respond to acute inflammation; however, little is understood about the dynamics and heterogeneity of these stress responses in HSPCs. Here, we performed single-cell sequencing of HSPCs during the sensing, response and recovery phases of the inflammatory response of HSPCs to treatment with the pro-inflammatory cytokine IFNα to investigate the HSPCs’ dynamic changes during acute inflammation. For the analysis of the resulting datasets, we developed a computational pipeline for single-cell time series. Using a semi-supervised response-pseudotime inference approach, we discover a variety of different gene responses of the HSPCs to the treatment. Interestingly, we were able to associate reduced myeloid differentiation programs in HSPCs with reduced myeloid progenitor and differentiated cells following IFNα treatment. Altogether, single-cell time series analysis have allowed us to unbiasedly study the heterogeneous and dynamic impact of IFNα on the HSPCs.

---

Inflammation is the body’s evolutionarily selected immune response to infection or tissue damage. It not only results in the activation and consumption of immune cells but is also accompanied by significant alterations in the function and output of hematopoietic stem and progenitor cells (HSPCs). Identifying how inflammatory stress regulates the fate of HSPCs and affects their function has become the subject of thorough scientific investigation in recent years (Caiado, Pietras, and Manz 2021). This started with our work and the work of others showing that pro-inflammatory cytokines such as interferons (IFNs) (Essers et al. 2009), (Baldridge et al. 2010), tumor necrosis factor-alpha (TNFα) (Pronk et al. 2011) or interleukin 1 (IL-1) (Pietras et al. 2016) are able to induce proliferation of normally quiescent hematopoietic stem cells (HSCs). Further investigations on for example, the mechanisms involved in stress-induced HSC activation or the response of progenitors have faced a significant challenge. Inflammation does not only impact the proliferation of hematopoietic cells, but it also induces extensive alterations in the expression of cell-surface proteins that are used as markers to distinguish different HSPC populations, with the strongest change being the increase in Sca-1 (Kanayama et al. 2020). This thus questions the reliability of using these surface markers in flow cytometry to identify and distinguish the different HSPC populations under inflammatory conditions. The recent development in single-cell expression profiling has advanced our understanding of HSPC heterogeneity (Watcham, Kucinski, and Gottgens 2019). Single-cell transcriptional profiles of HSPCs from these studies can now be used as reference datasets to identify individual HSPCs upon inflammation based on their transcriptional profiles, thus independent of cell-surface marker expression.

First single-cell experiments on HSPCs under inflammation have been reported; however, only a single time point was studied, thus providing only a snapshot of the response (Giladi et al. 2018). Here, we aimed to study the progression of inflammation-induced processes over time. This provides a unique opportunity to investigate the dynamics that underlie the response of the HSPC compartment to inflammation, but at the same time, generated computational challenges for the analysis of this type of single-cell dataset. Using a time series of single-cell RNA sequencing experiments covering the first 72 hours of the acute inflammation response *in vivo*, including the sensing, response, and recovery phase of HSPCs, we have worked out a computational pipeline to process and analyze single-cell time series. This pipeline includes cell type label transfer from the control time point to the treatment time points, characterization of the change in gene expression per cluster, visualization of the gene expression dynamics over pseudotime, and analysis of the cell type abundance over time. Using these approaches, we detected global and cluster-specific gene dynamics linked to different biological responses in HSPCs after IFNα treatment in our time series. We uncovered a reduction of myeloid progenitor cells associated with changes in transcriptional programs in multiple clusters. Thus, dynamic single-cell time series analysis has and will help us better understand how different cell types, genes, and processes change while the HSPC compartment progresses through the inflammatory response.

## Results

### A single-cell time series dataset capturing the dynamic inflammatory response of HPSCs

Biological responses such as acute inflammation are dynamic processes in which cells, tissues, and organisms undergo different phases of sensing differences, responding to these changes, and recovering upon successful response. Yet, often only single time points of these responses are investigated. Quiescent HSCs respond to inflammation by increased proliferation, which can be mimicked by treating mice with single pro-inflammatory cytokines, such as interferon alpha (IFNα). To gain a better understanding of the dynamics of the HSCs response to acute inflammation, we performed a time series experiment to cover the sensing, response, and recovery phases of the inflammatory response of HSCs to treatment with IFNα. While at 3 hours (3h) post-treatment, HSCs showed the first signs of sensing IFNα by increasing expression of interferon-stimulated genes (ISGs) (data not shown), only at 24 hours (24h) post-treatment HSCs reached a peak in increased proliferation (Fig. 1a) (Essers et al. 2009; Pietras et al. 2014). 72 hours (72h) post-treatment HSCs returned to quiescence (Fig. 1a). Unfortunately, inflammation does not only lead to increased proliferation of HSCs but is also accompanied by increased expression of several cell surface protein markers used to identify different cell types within the HSPC compartment. The most well-known example is the increase in the stem cell marker Sca-1 (Essers et al. 2009; Pietras et al. 2014; Kanayama et al. 2020). Using conventional marker-based flow cytometry including Sca-1, an increase in LSKs (Lin^-^ Sca-1^+^ cKit^+^) and a decrease in more committed LS^-^K (Lin^-^ Sca-1^-^ cKit^+^) progenitors is observed in response to inflammation (Fig. 1b-d). However, due to the increase in protein expression of Sca-1 (Fig. 1e,f) it is hard to predict whether these changes in frequency reflect an actual increase/decrease in cell frequency or are the result of contamination due to changes in protein expression of the cell markers in a given population or even both.

**Fig. 1.**
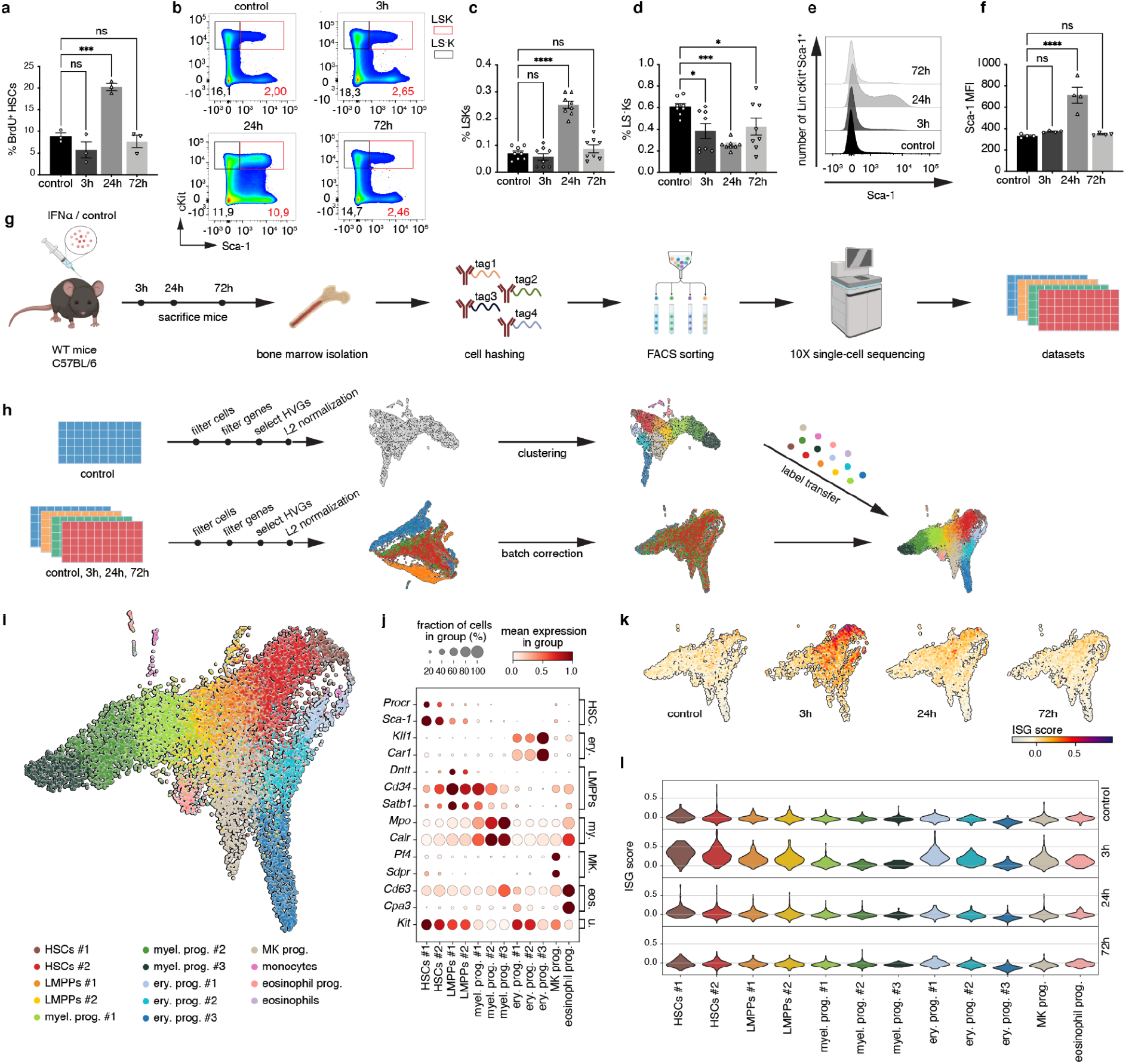
A single-cell time series RNA-seq dataset to characterize the response of HSPCs to IFNα treatment. **a,** Cell proliferation measured by 14 hour BrdU (18 mg/kg) uptake of HSCs (Lin^-^ Sca-1 ^+^ cKit^+^ CD150^+^ CD48^-^ CD34^-^) from control (PBS) or IFNα treated wild type (WT) mice (50000IU/20g mouse; 3h, 24h, and 72h). n=3 biological replicates. **b,** Representative FACS plots of Sca-1 and cKit expression on Lin^-^ bone marrow (BM) cells after control or 3h, 24h, and 72h time course IFNα treatment. **c,d,** Frequency of Lin^-^ Sca-1^+^ cKit^+^ (LSK) **(c)** and Lin^-^ Sca-1^-^ cKit^+^(LS^-^K) **(d)** cells in BM after control or 3h, 24h, and 72h time course IFN-α treatment. n=7 biological replicates. **e,f,** quantification (**e**) and statistical analysis (**f)** of Sca1 median fluorescence intensity (MFI) on Lin^-^ cKit^+^ cells. **g,** Scheme illustrating the experimental steps to acquire the single-cell time series RNA sequencing dataset. **h**, Scheme of the computational pipeline used to process the time series dataset. **i**, Two-dimensional UMAP embedding of cells from all time points, colored for the different identified clusters as indicated in the legend. **j**, Expression of marker genes in the different clusters in the control dataset. Erythroid (Ery.); Myeloid (My.); Eosinophils (Eos.); Universal (U.). **k**,**l**, UMAP embeddings (**k**), and violin plot (**l**) of the ISG score (see Methods) in the different cell clusters in the four different time points. Statistical significance in a, b, d, and e was determined by an ordinary one-way ANOVA using Holm-Šídák’s multiple comparisons test. At least 3 independent experiments were performed; *P≤0.05, ***P≤ 0.001 ****P<0.0001

To overcome these limitations, we adopted a single-cell RNA-seq approach to investigate the dynamics and heterogeneity of the stress response of stem and progenitor cells to IFNα treatment. Bone marrow cells were collected from IFNα-treated mice 3h, 24h, and 72h after treatment. Cells from PBS-treated mice were included as a control. LK (Lin^-^ and cKit^+^) cells were sorted to capture a wide spectrum of the HSPC transcriptional landscape. Since HSCs are much less frequent than the other populations in the LK gate, we enriched the sorted LK samples with LK CD150^+^ CD48^-^ CD34^-^ cells at a fixed ratio to the number of LK cells to guarantee sufficient numbers of stem cells for analysis (Supplementary Fig.1a). Inter-animal heterogeneity of the inflammatory stress response was addressed by performing cell hashing, for which cells from each biological replicate and time point were labeled with a unique hashtag antibody (Fig.1g and Methods). The cells from all four experimental time points (control, 3h, 24h, and 72h) were sequenced simultaneously. The resulting dataset was processed using a computational pipeline designed specifically for single-cell time series (Fig. 1h). First, the cells that did not meet the quality control standards (e.g. doublets, dying cells) were removed, resulting in a total count of 1600-3500 cells per time point (see Methods). Next, clustering of the cells identified 14 different clusters in the control subset. Using known marker genes for hematopoietic stem and progenitor cell types, 8 different cell types could be distinguished in the control subset (Fig. 1i,j). The assigned cell type labels were confirmed by scoring each cell for stemness (Supplementary Fig. 1b,c) and by comparing our dataset to a previously published dataset (Supplementary Fig. 1d) (Nestorowa et al. 2016). The cell type labels in the control subset were transferred to the other three treatment time points (Fig. 1h and Methods). Marker gene expression confirmed the cell type labels in the response time points (Supplementary Fig. 1e). Analysis of the hashtags showed that the biological replicates in each time point had comparable abundances of each cell type label (Supplementary Fig. 1f). In addition, expression analysis of the interferon α/β receptor (*Ifnar1* and *Ifnar2*) confirmed that all clusters expressed the interferon α/β receptor and thus were able to directly respond to IFNα (Supplementary Fig. 1g,h). As a first measure of the inflammatory response, the expression of interferon-stimulated genes (ISGs) was scored (Fig. 1k,l). At 3h all clusters underwent a change in their ISG expression, indicating that the whole HSPC compartment sensed the IFNα treatment (Fig. 1k). However, the data also indicate great heterogeneity in the ISG response between and within clusters with HSC clusters showing the biggest change (Fig. 1l).

### Inflammation response is defined by global and cluster-specific changes in gene expression

To gain a comprehensive understanding of all genes that characterize the IFNα response, we next performed differential gene expression analysis. Differentially expressed genes were selected between the control subset and every treatment time point individually to get a set of response genes from any stage of the response (see Methods). The analysis identified a total of 2501 significant response genes. Expression profiles of the response genes showed that the expression of some genes changed globally (e.g. *Cox7c*), whereas the expression of others was more specifically changed in few or one cluster (e.g. *Sec61g* and *Mnda*) (Fig. 2a). To investigate in which cluster(s) genes were changing the most, the top 500 most significant response genes were scored for the total expression change in each cluster (change score; see Methods). After calculating the total change for each cluster, the response genes were categorized into 14 different groups using hierarchical clustering (Fig. 2b), confirming a wide variety of globally responding genes (groups 1-5), as well as cluster-specific responding genes (groups 6-14). To exclude that the differences between clusters were a result of completely different expression profiles, the similarity between a cluster’s expression profile and the expression profile in the whole dataset was calculated (Supplementary Fig. 2a). The results indicate that most expression profiles follow similar patterns in all clusters.

**Fig. 2.**
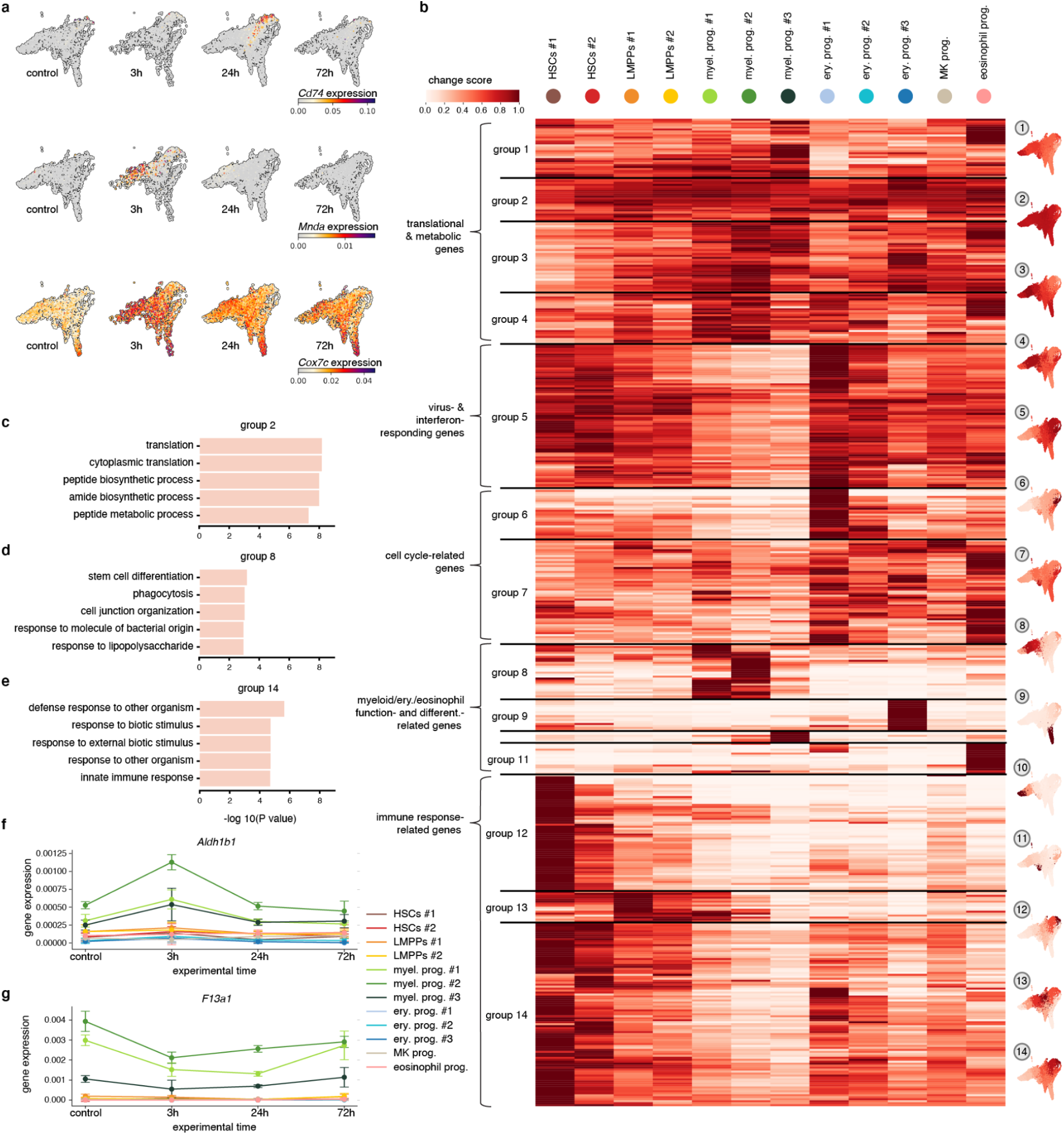
Inter-cluster analysis of response genes shows both global and cluster-specific responding genes. **a,** UMAP embeddings with the expression of response genes *Cd74, Mnda*, and *Cox7c* in control and 3h, 24h, and 72h post IFNα treatment. **b,** Change score (see Methods) in each cluster for the top 500 response genes, grouped using hierarchical clustering. UMAPs on the right show the expression change for each cell cluster averaged over all response genes in the corresponding group. On the left are terms summarizing the functional annotation of the response genes associated with the groups. **c,d,e** GO terms associated with group 2 (**c**), 8(**d**), and 14(**e**). The length of each bar represents the statistical significance of each term. **f,g,** Mean expression of *Aldh1b1* (**f**), and *F13a1* (**g**) in each cluster over time.

Gene ontology (GO) enrichment analysis (Fig. 3c-e and Supplementary Fig. 2b-l) of the global response gene signatures in groups 1-4 revealed an overrepresentation of terms associated with translation and metabolism (Fig. 2c and Supplementary Fig. 2b-d). This is in line with reports of HSPCs undergoing massive changes in the metabolism under inflammatory stress (Karigane and Takubo 2017). In addition, global response genes from group 5 (Supplementary Fig. 2e) showed enrichment for categories associated with immune response and response to type-I interferon, further supporting the ISG expression data (Fig. 1k), indicating that all cells sense the changes in IFNα levels. Interestingly, expression changes in the HSC-enriched groups 12 and 14 were also associated with immune response and response to type-I interferon (Fig. 2e and Supplementary Fig. 2k). However, these changes were different from the changes in group 5, suggesting an HSC-specific immune response, which is different from the immune response in progenitors. HSC-enriched groups 12 and 14 also included GO terms such as regulation of T cell activation and antigen processing and presentation (Fig. 2e and Supplementary Fig. 2k), which correspond to the newly identified role of HSCs as immunomodulators (Hernández-Malmierca et al. 2022), which would be strengthened under inflammation. Besides global and HSC-specific response genes related to immune response, change score analysis also identified groups of response genes enriched in committed progenitors related to progenitor-specific processes. For example, erythroid and eosinophil progenitor-enriched groups 9 and 11 showed an overrepresentation of processes related to erythrocyte differentiation terms and myeloid development (Supplementary Fig. 2h,j). Change scores in groups 8 and 10 were largest for myeloid progenitors and connected with biological processes such as phagocytosis, myeloid leukocyte-mediated immunity, as well as stem cell differentiation, which are characteristic functions of this cell type (Fig. 2d and Supplementary Fig. 2i). Thus, with the change score analysis both global and cluster-specific signatures were identified. The analysis highlights that HSCs are the major responders to inflammation in the HSPC compartment and both global and HSC-specific inflammation signatures are present, indicating heterogeneity in the inflammatory response between the clusters.

**Fig. 3.**
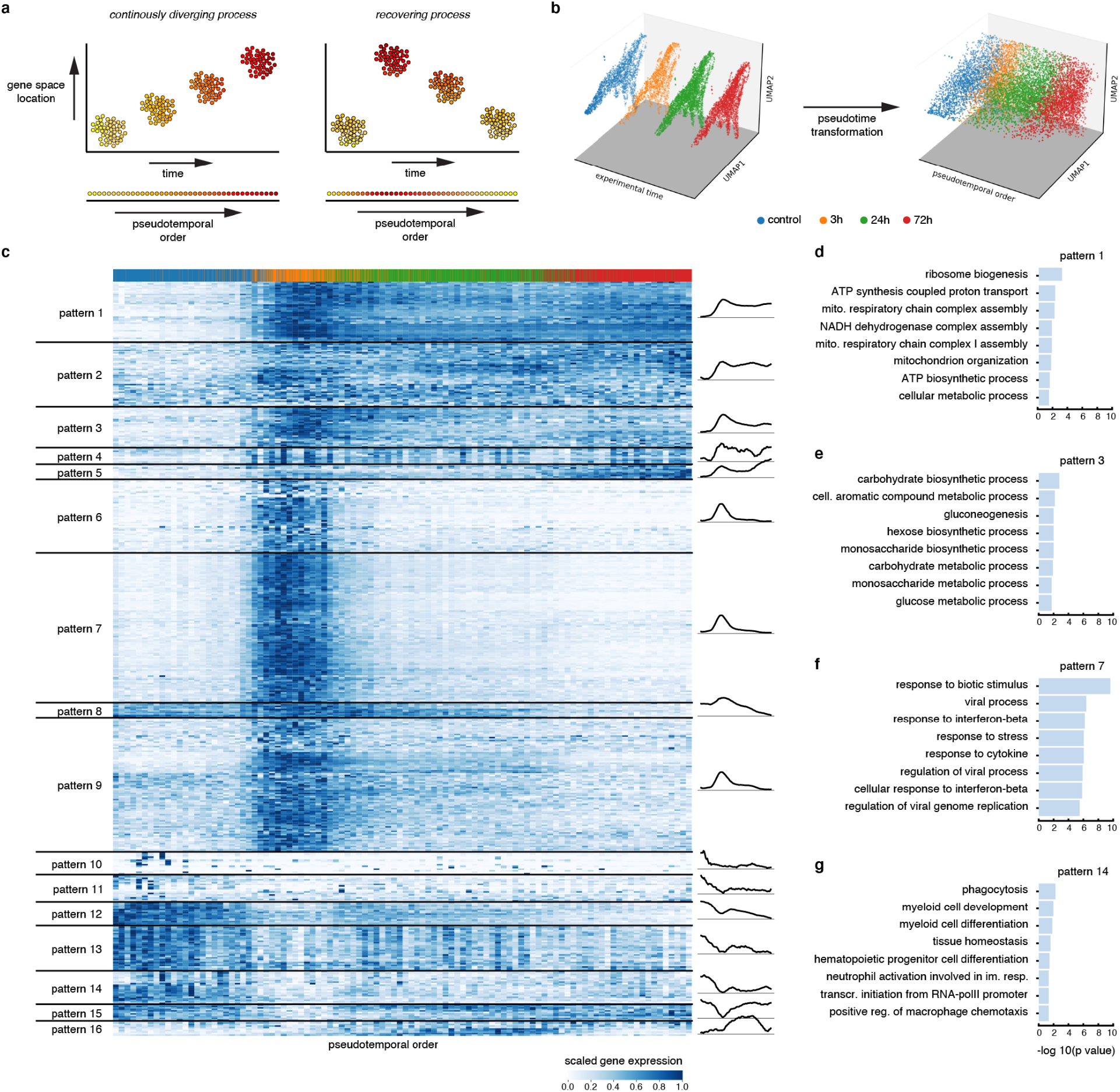
Implementing response pseudotime to characterize the dynamics of gene expression in response to IFNα treatment. **a,** Illustration of the difference between time series that capture continuously diverging processes (such as development), and recovering processes (such as acute stimulation). **b,** Three-dimensional embedding of the HSPC dataset with experimental timepoints (left) or pseudotemporal ordering (right) on the z-axis (x- and y-axis: UMAP). **c,** (smoothed) Expression of the top 500 response genes, with cells ordered by pseudotime and genes grouped by pattern using hierarchical clustering. A graphical representation of the mean pattern in each pattern group is shown on the right. **d,e,f,g** GO terms associated with gene patterns 1 (**d**), 3 (**e**), 7 (**f**), and 14 (**g**). The length of each bar represents the statistical significance of each term.

### The pseudotemporal ordering of cells enhances the resolution of gene dynamics

The change score analysis gave an overview of all response genes without taking into account the dynamics of the response. Therefore, in the next step, the expression dynamics of the response genes were explored. When zooming into the expression of individual genes in time, different expression patterns were observed for genes within the same group. For example, the responses of *F13a1* and *Aldh1b1* were both assigned to group 8. However, the temporal expression dynamics differed between both genes; *F13a1* expression steadily went down (Fig. 2g), whereas *Aldh1b1* was upregulated with a peak at 3h and a full recovery to original (control) expression levels at 24h (Fig. 2f). To improve the characterization of the expression patterns, we wanted to leverage the single-cell resolution of our dataset. Therefore, we aimed to construct a pseudotime axis in the gene expression space to describe the inflammatory response. In datasets covering a developmental process or disease progression, the asynchrony of cells can be leveraged to infer a pseudotemporal ordering of cells (using methods such as diffusion pseudotime (Haghverdi et al. 2016), Monocle (Qiu et al. 2017), etc.). These methods are generally based on cell neighborhood relations, with the assumption that the further apart (in Euclidean, diffusion space, etc.) two cell states are, the longer the typical transition time between them, hence longer pseudotime (Fig. 3a). However, this assumption is violated for the type of post-drug treatment time series data we have here, where the largest transcriptional change is observed shortly after stimulation but (presumably) diminishes as cells relax to a more control-like state over a longer time (Fig. 3a). Cells from different experimental time points generally appear either completely intermingled or completely disconnected from one another when viewed over the first principal components (Supplementary Fig. 3a). This renders the unsupervised, distance-based pseudotime methods unsuitable for detecting the temporal order of response dynamics. Thus, to get a higher temporal resolution of the response progression from the four time points, we used a new approach that finds a pseudotime axis that best correlates with the cells experimental time labels. We find a transformation of the cells’ expression matrix that reconstructs the experimental time labels plus an error term that we try to minimize (Supplementary Fig. 3b and Methods). In essence, this approach assigns a positive/negative weight to each gene (depending on its correlation or anticorrelation with the experimental time labels), in such a way that the resulting weighted sum of the gene expressions approximates the experimental time labels of the cells best. The genes that positively correlate with experimental time turn out to be often associated with the immune response, whereas anticorrelated genes display enrichment in processes such as translation (Supplementary Fig. 3c-g). Using the pseudotemporal order of the cells, the different expression dynamic patterns that follow IFNα treatment were explored. Response genes were categorized into 16 patterns based on their pseudotemporal dynamics using hierarchical clustering (Fig. 3c and Supplementary Fig. 3h). Each pattern represents a cluster of genes with similar expression dynamics following IFNα injection, which can be roughly subdivided in upregulation (patterns 1-9 and 16) and downregulation (pattern 10-15) patterns. The patterns showed a broad diversity in the speed of sensing, response and recovery. A few patterns (patterns 5,8,12) do not reach a plateau until the latest pseudotime points and imply an ongoing trend of change. Genes in patterns 6, 7, and 9 show quick sensing, response, and recovery associated with immune, interferon, and viral processes (Fig. 3f). Other patterns resembled a similar fast increase but with slower recovery, as seen for pattern 3, which is enriched for metabolic processes (Fig. 3e). In addition, the heatmap in figure 3c showcases a variety of gene dynamics that were different from the rapid response and recovery IFN-response (patterns 6,7 and 9), such as a sustained upregulation in pattern 1, which was associated with translation and other biosynthetic processes (Fig. 3d). In contrast, several gene patterns encompassed genes that were downregulated (gene patterns 10-14). After the initial decrease in expression many of these genes failed to recover to initial expression levels. The majority of these genes were linked to myeloid development and differentiation (Fig. 3g). This would suggest alterations in myeloid differentiation upon the IFNα treatment, an observation described for many other pro-inflammatory cytokines (Matatall et al. 2014; Pietras et al. 2016; Yamashita and Passegué 2019) but IFNα.

### Response pseudotime reveals a landscape of gene dynamics in HSPCs following IFNα treatment

To decipher the dynamic changes in the inflammatory response in the different clusters, we combined the information on how response genes changed their expression (Fig. 3c) with whether these changes were global or cluster-specific (Fig. 2b). The result condenses the plenitude of information in the complete singlecell time series into a single visualization, which considerably eases the search for (groups of) biologically relevant genes (Fig. 4a).

**Fig. 4.**
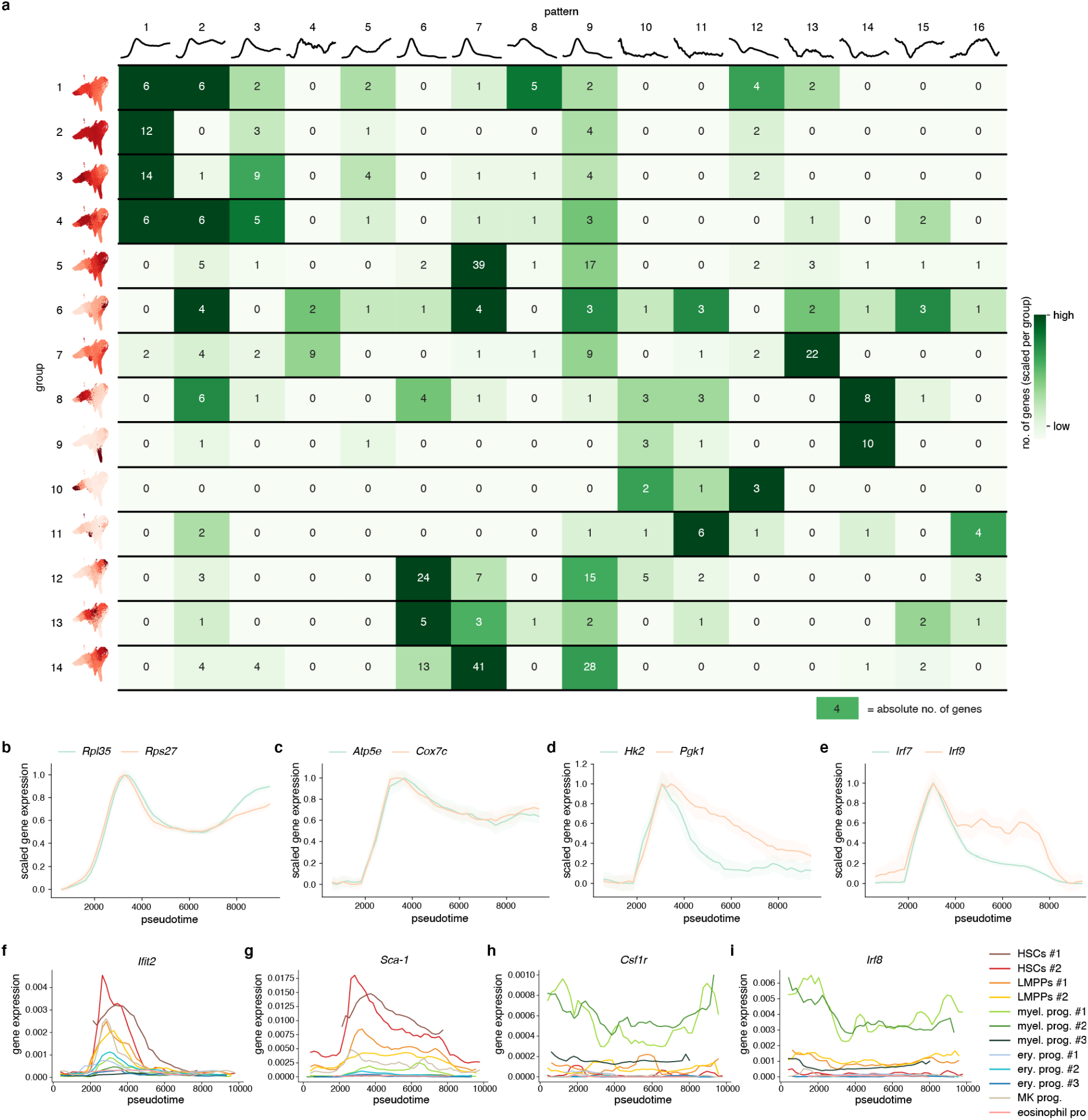
Response pseudotime reveals a landscape of gene dynamics in HSPCs following IFNα treatment. **a,** Visual summary of the HSPC time series showing the breakdown of the response gene patterns in each change score group. The numbers in each cell represent the absolute number of genes (e.g. 5 response genes in change score group 1 display pattern 1). The colors represent the number of genes scaled for each change score group. **b-e,** Examples of gene expression in pseudotime for translation (*Rpl35, Rps27*) (**b**), metabolism (*Atp5e, Cox7c, Hk2, Pgk1*) (**c,d**), and inflammation (*Irf7, Irf9*) (**e**) specific genes. **f-i,** Pseudotemporal expression of HSC-specific genes (*Ifit2, Sca-1*) (**f,g**) and myeloid-specific genes (*Csf1r, Irf8*) (**h,i**) in different clusters.

The two most commonly found patterns among the groups are patterns 2 and 9, both showing a fast increase combined with a slow (pattern 2) or fast (pattern 9) recovery. Genes from these patterns were mainly related to immune responses, highlighting that the sensing and response to IFNα is fast and present in all groups and clusters. The slower recovery rate in pattern 2, however, also indicates that even the general immune response does not show full recovery within 72h.

Interestingly, other patterns of fast sensing and response followed by sustained upregulation (patterns 1 and 3) were enriched in the groups with a global signature (groups 1, 2, 3, and 4). Several of these genes were ribosomal (e.g., *Rpl35* and *Rps27;* Fig. 4b), suggesting that biosynthetic activity increased in most HSPCs early in the treatment and remained active even in the recovering phase. In addition, several genes following the sustained upregulation pattern were metabolic. Oxidative phosphorylation (OXPHOS) genes and mitochondrial enzymes (e.g., *Atp5e* and *Cox7c;* Fig. 4c) showed these patterns of prolonged upregulation. On the other hand, glycolytic genes *Hk2* and *Pgk1* showed a quick response and recovery (Fig. 4d). Thus, in contrast to previous reports suggesting a binary (on/off) switch between glycolysis and OXPHOS (Suda, Takubo, and Semenza 2011), our data suggests that an initial upregulation of glycolytic and a sustained upregulation of OXPHOS genes go hand in hand in inflammation responding HSPCs.

In contrast to the heterogeneity in dynamics observed globally, HSC-enriched groups (groups 5, 12, and 14) are mainly enriched with gene patterns that increase very early after treatment and quickly return to homeostatic levels (patterns 6, 7, and 9). Examples of such genes are *Irf7* and *Irf9* in group 5 (Fig. 4e) or *Ifit2* and *Sca-1* in groups 12 and 14 (Fig. 4f,g and Supplementary Fig. 4a,b, which indicates the confidence intervals for the expression of these genes). Thus, the majority of HSC-enriched groups follow rapid sensing, responding, and recovery dynamics, with the majority of gene changes preceding the peak in proliferation response in these cells. In addition, the majority of HSC-enriched dynamic changes are within gene groups linked to interferon and immune response, again highlighting the specific, fast HSC-specific immune response.

In contrast to HSC-specific groups, committed progenitor-specific groups (8, 9, 10, and 11) were strongly enriched in genes that exhibited persistent downregulation (patterns 10, 11, 12, 14). In the myeloid progenitor-specific groups 8 and 10, many of these genes were associated with myeloid cell differentiation and functional programs e.g., Csf1r; Irf8 (Fig. 4h,i and Supplementary Fig. 4c,d), suggesting reduced myeloid differentiation in the myeloid progenitor clusters.

In summary, response pseudotime has shed light on the heterogeneity in gene dynamics in the HSPC compartment during the induction of inflammation. Whereas global groups encompass diversity in gene patterns, cluster-enriched gene groups show far less variation and more specificity, with HSCs being the fast responders and recoverers, whereas committed myeloid progenitors showing sustained downregulation of genes.

### Single-cell abundance analysis shows myeloid depletion and HSC enrichment following IFNα treatment

To investigate whether reduced transcriptional programs for myeloid differentiation and function upon IFNα treatment also impacted the size of the progenitor compartment, we performed differential abundance analysis. We applied the Milo algorithm (Dann et al., 2021), which models cellular states as overlapping neighborhoods on a KNN graph, rather than relying on clustering cells into discrete groups (see Methods). At a false discovery rate (FDR) of 10%, we could observe multiple neighborhoods that were differentially abundant (Fig. 5a). Neighborhoods received a cell-type label based on the most predominant cluster in the neighborhood. Even though most progenitor-enriched clusters showed a reduction at 3h, the majority returned to normal by 24h or 72h, except for the most differentiated myeloid progenitors (Myel. prog. #3), which were sustainably reduced, even at 72h posttreatment (Fig. 5b,c). The abundance of HSCs only slightly increased in HSCs #2 at 3h, but was back to normal at 72h (Fig. 5b,c). Thus, this unbiased (i.e., abundance analysis of cell types based on the expression of several genes rather than specific markers) single-cell investigation of the cell type frequency in the HSPC compartment showed that acute IFNα treatment resulted in a slight enrichment of HSCs at the early time point, but a sustained reduction in the most committed myeloid progenitors over the whole time course of the response. This is in contrast to the current notion in the field claiming that the decreased frequency of LS^-^K (comprising myeloid, erythroid, and megakaryocytic progenitors) and concurrent increase in LSKs (comprising HSCs and LMPPs) upon IFNα stimulation (Fig. 1b-e) is mainly the result of contaminating myeloid progenitors that have reacquired *Sca-1* expression (and would fall into the LSK gate) (Pietras et al. 2014; Kanayama et al. 2020). However, when analyzing *Sca-1* gene expression in our data set, no change in *Sca-1* mRNA transcripts could be observed in myeloid progenitors (Fig. 5d,e). In contrast, IFNα induced upregulation of *Sca-1* was solely observed in HSC and LMPP clusters, with the strongest increase in the HSCs (Fig. 5d,e). Hence, this unbiased investigation of the different clusters based solely on their gene expression identifies a true change in myeloid population size and not a shift in populations due to marker change.

**Fig. 5.**
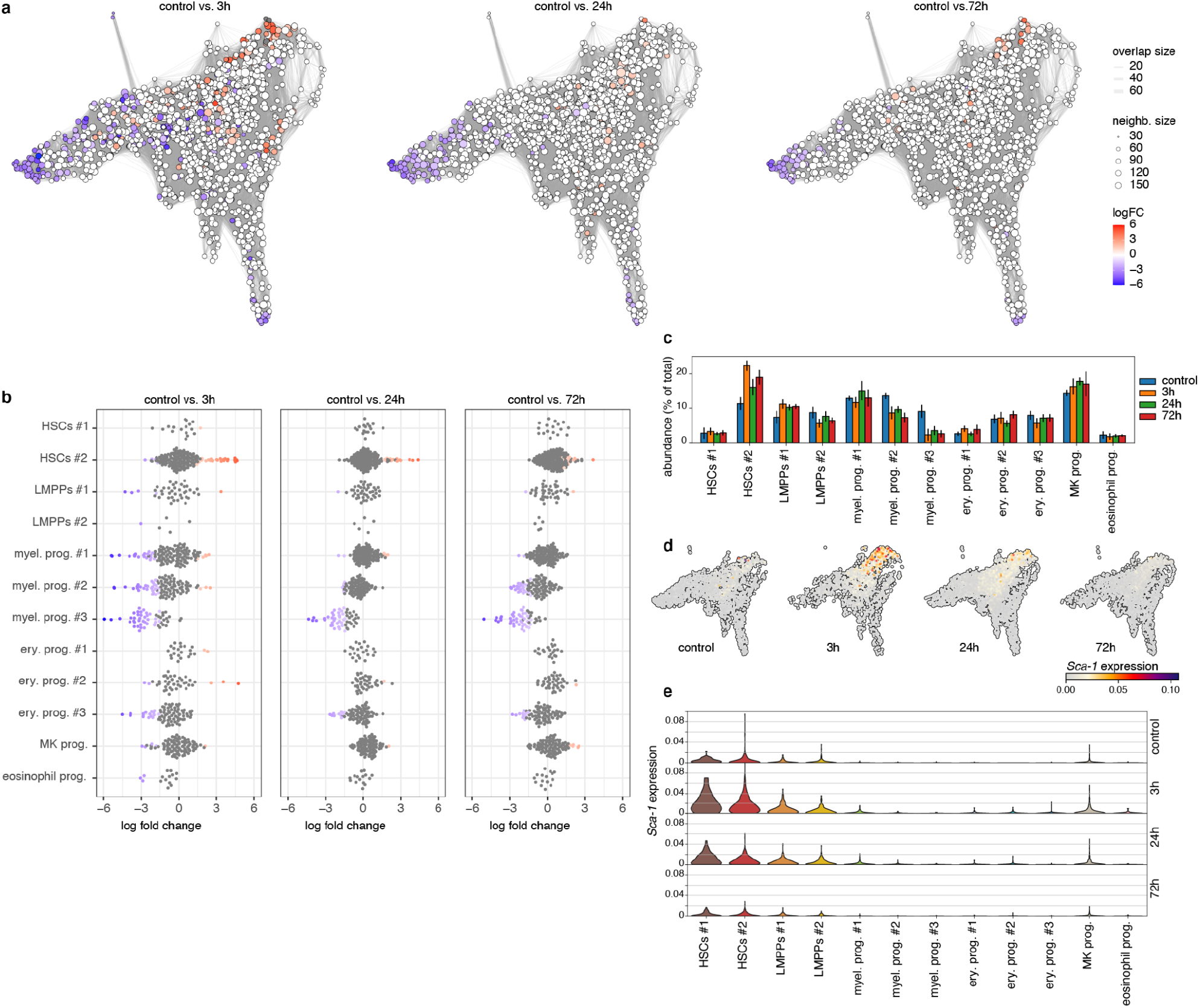
Abundance analysis reveals a sustained reduction in myeloid progenitors following IFNα treatment. **a,** Neighborhood graphs with the results from Milo differential abundance testing between the control dataset and post IFNα treatment subsets (3h, 24h, 72h). Nodes represent neighborhoods, coloured by their log fold change (red: more abundant, blue: less abundant, white: non-differentially abundant.) Graph edges represent the number of cells shared between two neighborhoods. **b,** Beeswarm plot of the distribution of log fold changes in each cluster. Neighborhoods are assigned to clusters based on the most commonly found cluster label in the neighborhood. **c,** Relative abundance of clusters in each time point. **d,e,** UMAP embeddings **(d)** and violin plot **(e)** of *Sca-1* expression in the control, 3h, 24h, and 72h timepoints.

### Myeloid depletion coincides with changes in transcriptional programs

The reduction in the abundance of the myeloid progenitors could be caused by an increased egress of these cells from the bone marrow, a loss of these cells due to cell death, or a reduction in differentiation towards myeloid progenitors. Myeloid progenitors were not observed in the blood at any time point following IFNα treatment (data not shown). However, the number of myeloid progenitors in the spleen decreased at 24h, in line with the reduction observed in the bone marrow (Supplementary Fig. 5a). This suggests that the reduced abundance of the myeloid progenitors in the bone marrow is not due to increased egress into the blood or spleen. To investigate whether reduced levels of myeloid progenitors were the result of increased cell death, gene patterns of pro-survival genes (*Bcl2, Birc2*, and *Birc5*) were analyzed and found to be decreased following IFNα treatment (Fig. 6a). Yet in *Bax*^-/-^*Bak*^-/-^ double-knockout mice, in which cells are unable to undergo apoptosis do to the loss of the pro-apoptotic proteins *Bax* and *Bak*, a similar reduction of myeloid progenitors was observed as in wild type mice (Fig. 6b), indicating that apoptosis was not the reason for the reduction in myeloid progenitors. However, this result does not exclude the involvement of other forms of cell death, like necroptosis and pyroptosis. Necroptosis gene expression was not altered in any of the cell types (Fig. 6c and Supplementary Fig. 5b-e), but an expression of pyroptosis genes, e.g., *Casp1* and *Casp4*, was increased, suggesting an early increase in pyroptosis in the myeloid progenitors in response to the treatment, possibly resulting in a reduction of these cells (Fig. 6c,d and Supplementary Fig. 5b-d,f).

**Fig. 6.**
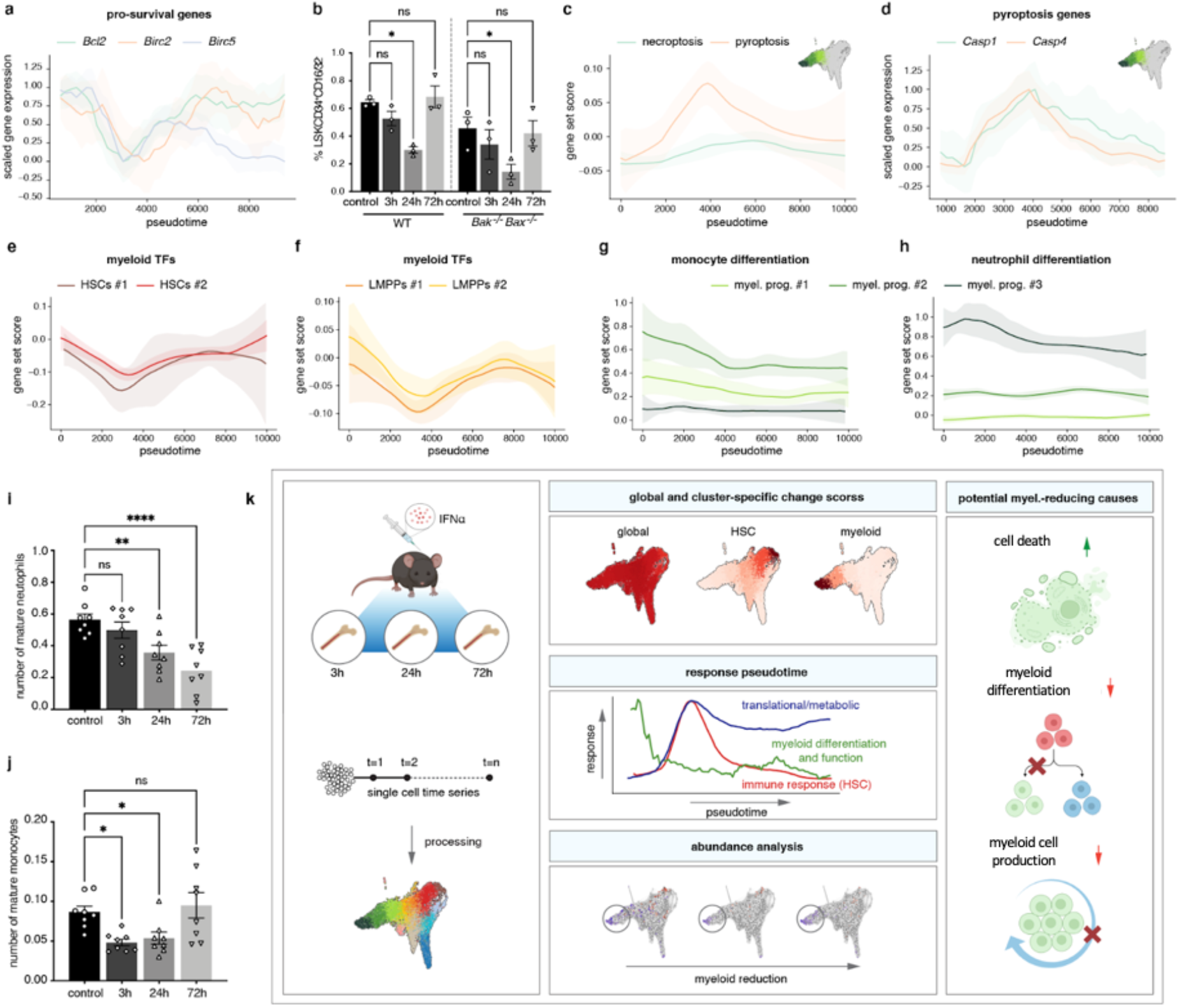
Reduced myeloid differentiation bias and increased cell death signature partially explain myeloid population’s reduction upon inflammation. **a,** Pseudotemporal expression of pro-survival genes (*Bcl2, Birc2, Birc5*). **b,** Flow cytometric analysis of BM frequency of myeloid progenitors (Lin^-^ Sca-1^-^ cKit^+^ CD34^+^ CD16/32^+^) following IFNα treatment in WT and *Bax*^-/-^*Bak*^-/-^ double knockout mice at 3h, 24h, and 72h following IFNα or control (PBS) treatment. n= 3 biological replicates. **c,** Score of necroptosis gene signature and pyroptosis gene signature in all myeloid progenitors plotted in pseudotime. **d,** Pseudotemporal expression of two pyroptosis genes (*Casp1, Casp4*) in all myeloid progenitors **e, f,** Score of (murine) myeloid transcription factors in HSCs **(e)**, and LMPPs **(f)** plotted in pseudotime. **g,h,** Score of monocyte **(g)** and neutrophil **(h)** differentiation genes in all three myeloid progenitor clusters plotted in pseudotime. **i,j,** Flow cytometric analysis of blood neutrophils (B220^-^ CD4^-^ CD8^-^ Ly6G^+^ CD11b^+^) **(i)** and monocytes (B220^-^ CD4^-^ CD8^-^ Ly6G^-^ CD11b^+^ CD11c^-^ F4/80^-^) **(j)** normalized to the whole blood leukocyte count as measured by hemavet at 3h, 24h, and 72h injection of IFNα or control (PBS) treatment in WT mice. n= 8 biological replicates. **k.** Graphical abstract of the paper. Statistical significance in **i,j** was determined by an ordinary one-way ANOVA using Holm-Šídák’s multiple comparisons test. At least two independent experiments were performed; *P≤0.05,**P≤0.01, ***P≤ 0.001, ****P<0.0001. Data represent mean ± standard error of the mean (SEM).

To investigate whether insufficient production of new cells might also play a role in the sustained depletion of myeloid progenitors, the expression of genes involved in myeloid lineage priming was analyzed. By calculating a gene score for known myeloid transcription factors that have been reported to impact both stem and progenitor cells, a downregulation in myeloid priming in HSCs and the more differentiated LMPPs was present in the early stages of the IFN response (Fig. 6e,f). Additionally, a reduction in cell cycle and purine nucleotide synthesis genes was observed in myeloid progenitors, suggesting that both myeloid differentiation in HSCs and LMPPs as well as cell production in myeloid progenitors, is affected (Supplementary Fig. 5g-i). We also tested if the post-inflammation decline in differentiation bias towards myeloid progenitors could be captured by RNA-velocity (La Manno et al. 2018) of the cells based on the unspliced versus spliced reads from several genes. However, computing the cell velocities for each of the datasets in the time series was hindered by the yet insufficient robustness of current velocity estimation approaches as well as the poor quantification of unspliced versus spliced mRNA, especially in the 3’-end gene expression short reads captured by the 10X sequencing platform (Supplementary Note 1).

Along with the reduction in myeloid production related genes, neutrophil, and monocyte transcriptional signatures were also downregulated in myeloid progenitors over the pseudotime axis (neutrophil: myel. prog. #1; monocyte: myel. prog. #3) (Fig. 6g,h), suggesting continuously reduced differentiation of myeloid progenitors to mature myeloid cells. Indeed, the number of neutrophils in the blood gradually decreased over time (Fig. 6i). Monocyte numbers showed recovery at 72h after an initial reduction (Fig. 6j). Taken together, these *in vivo* cell analyses and gene expression data indicate an increase in pyroptosis signature combined with reduced myeloid differentiation both at the stem cell level, as well as in committed progenitors, and a reduction in the cell production machinery in progenitors, resulting in reduced myeloid progenitors in the bone marrow, and lower level of mature myeloid cells in the blood (Fig. 6k).

## Discussion

Analysis of single-cell RNA-seq time series is nontrivial because of its high complexity, regarding the inclusion of multiple cell types, a high number of genes, and the extra dimension of (pseudo-) time. However, these types of experiments allow for marker-independent, unbiased analysis of dynamic responses of heterogeneous cell populations, such as the response of the HSPC compartment to inflammatory stress. To overcome the challenges of analyzing such datasets, we designed a computational pipeline for processing and analyzing single-cell RNA-seq time series, in which clusters were labeled based on the expression of multiple (n = 3326) genes after correcting for treatment effects (Fig. 6k), thus avoiding relabeling of the cells undergoing inflammation as separate populations. This will ensure the study of the same cell type over time. Hence, cell type identity is reliably retrieved, even though the conventional marker genes might be subject to changes, as is the case during inflammation.

Furthermore, we designed measures that make the temporal dynamics more comprehensible and provide a (visual) entry point into all the information in the data. Unsupervised methods that are merely based on cell-to-cell proximities, are unable to capture the pseudotemporal order of the cells in our data, where the cell states after relaxation become more similar to the cell states before treatment. Therefore, we implemented a semi-supervised, i.e., using the experimental time labels information, method for inference of response pseudotime. Using a minimal linear regression model, our response pseudotime reconstruction enabled capturing the fine-grained expression changes and dynamical patterns beyond the four discrete experimental time points (Fig. 6k). Recently, alternative semi-supervised methods such as psupertime (Macnair, Gupta, and Claassen 2022), or methods suitable for other temporal patterns (e.g., periodic dynamics) that linear matrix transformations may not capture are also being investigated.

Although many studies have investigated the role of inflammation on HSC function, changes in marker expression on these cells have made it challenging to examine the impact of inflammation on the heterogeneity and molecular changes over time in the HSPC compartment. Our response pseudotime approach allowed us to highlight the dynamic nature of HSPCs’ response to IFNα with global and cell typespecific distinct molecular patterns of gene expression and biological processes (Fig. 6k). Whereas global gene groups and patterns were heterogeneous in dynamics and linked to diverse processes such as metabolism, translation, and inflammation, HSC-specific gene groups and patterns followed rapid sensing, response, and recovery and were enriched for distinct inflammation-related genes, suggesting global as well as stem cell-specific responses to inflammation. Interestingly, several gene groups did not recover from their induced increased or decreased expression along the response pseudotime of 72 hours posttreatment, suggesting ongoing cascades of molecular changes or possibly also irreversible changes leaving a mark in the activated cells. This would be in line with a recent study showing that HSC function is irreversibly attenuated by temporally discrete inflammatory events (Bogeska et al. 2022).

Emergency myelopoiesis, i.e., increased production of myeloid cells, has been described in response to many pro-inflammatory cytokines and infections (Manz and Boettcher 2014). However, thus far, we have not been able to identify any impact on myeloid production or differentiation upon IFNα treatment due to extensive changes in stem cell-specific marker expression upon inflammation (Demerdash et al. 2021). Others claimed that decreased frequency of myeloid progenitors and the simultaneous increase in LSKs upon *in vitro* treatment of HSPCs with IFNα was mainly the result of myeloid progenitors reacquiring Sca-1 expression (Pietras et al. 2014; Kanayama et al. 2020). With our unbiased investigation of the different clusters, defined solely by their gene expression, we could now show that IFNα-induced *Sca-1* gene expression only occurred in immature HSCs and LMPPs (Fig. 5d). Even though CITEseq analysis of HSPCs should be performed to confirm these results at the Sca-1 protein level, our data does indicate that the LSK expansion observed in flow cytometry is mainly due to the enrichment of the HSCs and multipotent progenitors and not myeloid progenitor populations shifting into the LSK gate.

Unlike other proinflammatory cytokines such as TNFα (Yamashita and Passegué 2019) and IL1β (Pietras et al. 2016), we did not find characteristics typical of emergency myelopoiesis. Instead, abundance analysis showed a decrease in myeloid progenitor numbers; gene expression related to myeloid priming was downregulated in all clusters from immature HSCs to committed myeloid progenitors; and myeloid-derived mature neutrophils were continuously reduced in the blood (Fig. 6k). In addition, response pseudotime analysis revealed changes in expression of genes related to pyroptosis, suggesting that reduced levels of myeloid progenitors might be the result of a combination of impaired myeloid differentiation with increased cell death via pyroptosis. Interestingly, upon infection with Mycobacterium tuberculosis, HSCs are reprogrammed to limit their commitment towards myelopoiesis via a type I IFN signaling axis (Khan et al. 2020). In this same study, they showed that IFNα induces RIPK3-mediated necroptosis in myeloid progenitors. However, RIPK3 is a component of both pyroptosis and necroptosis depending on other proteins participating in these pathways (Shlomovitz, Zargrian, and Gerlic 2017). Differentiation and cell death pathways are not only regulated at the transcriptional level. Thus post-translational analysis and additional functional approaches need to be performed to unravel further the programs controlling the IFNα-induced reduction in the myeloid progenitors in the bone marrow. Thus, our time course data suggest an unanticipated impact of IFNα on the differentiation and production of myeloid cells, highlighting the diverse impact of the same pro-inflammatory agonist on related but distinct cell types at different timepoints in the response. This link between IFNα and reduced production and levels of myeloid cells such as neutrophils not only helps us to better understand the impact of inflammation on the whole hematopoietic compartment. It will also help to understand better the role of IFNα in disease settings such as the autoimmune disease systemic lupus erythematosus (SLE) in which neutrophil dysfunction plays an integral role in disease pathogenesis (Kaplan 2011) and IFNα is associated with adverse outcomes (Rönnblom and Leonard 2019).

## Methods

### Mouse models

All animal experiments were approved by the local Animal Care and Use Committees of the German Regierungspräsidium Karlsruhe für Tierschutz und Arzneimittelüberwachung (Karlsruhe, Germany). Mice were kept under specific pathogen-free conditions (SPF) in ventilated cages (ICV) in the animal facility of the German Cancer Research Center (DKFZ). Mice used for experiments were between 10-20 weeks old at the beginning of the respective experiments. *SclCreERT bax*^-/-^*bak*^-/-^ mice were on a C57BI/6 background (Takeuchi et al. 2005), and treated for 5 days with 2mg/day tamoxifen. IFNα treatment was started 4 weeks post tamoxifen treatment. C57Bl/6 (WT) mice were bred at the DKFZ animal facility or bought from JANIVER lab. Mice were sacrificed by cervical dislocation according to German guidelines.

### IFNα treatment of mice

Mice were injected subcutaneously with 50.000 international units (IU) of recombinant mouse IFNα per 20g mouse (Milteny Biotech). Recombinant mouse IFNα was diluted in PBS and control mice were injected with 100 μl PBS.

### Isolation of bone marrow (BM), spleen, and blood for flow cytometry analysis

Blood was collected from the vena facialis by sub-mandibular bleeding into EDTA-coated collection tubes. Blood was either analyzed automatically with a Hemavet cell counter (Drew Scientific) or stained for flow cytometry after initial RBC lysis by incubation with ACK lysis buffer for 20 mins. Cells were stained for Ter119, CD4, CD8, CD11b, CD11c, Ly6G, CD41, B220, F4/80. BM cells were isolated from the femur, tibia, hip bone, and spine by bone-crushing. Splenocytes single-cell suspension was obtained by mashing the spleen through a 40 μm EASYstrainer™ (greiner bio-one). After ACK lysis BM cells and splenocytes were stained using antibodies for CD117 (cKit), Sca-1, CD150, CD48, CD34, CD16/32, and lineage antibodies (CD4, CD8, CD11b, Gr-1, B220, and Ter119). For the BrdU incorporation assay, BrdU (18mg/kg, Sigma-Aldrich) was administered i.p. for 14 hours prior to harvesting the BM. The BD Pharmingen™ BrdU Flow Kit protocol was used to stain for BrdU. For flow cytometry analysis the LSR Fortessa or LSRII were used (BD Biosciences). Flow data were analyzed using BD FACS DIVA v8.0.1 and Flowjo (v10).

### FACS sorting

For FACS sorting of single cells, BM cells were isolated and RBC lysed as described above. This was followed by lineage depletion using a lineage antibody cocktail against CD4, CD8, CD11b, B220, Gr-1, and Ter119 and incubation with Dynabeads^®^ Magnetic Beads (Invitrogen). Lineage-depleted BM cells were stained with Zombie Yellow viability dye (BioLegend) followed by incubation with the following antibodies: CD117, Sca1, CD150, CD48, CD34, and lineage antibodies (CD4, CD8, CD11b, Gr-1, B220, and Ter119) together with one of the hash antibodies (TotalSeq™-A0301 anti-mouse Hashtag 1 Antibody, TotalSeq™-A0302 anti-mouse Hashtag 2 Antibody, TotalSeq™-A0303 anti-mouse Hashtag 3 Antibody, TotalSeq™-A0304 anti-mouse Hashtag 4 Antibody) (BioLegend, TotalseqA antibodies). The 4 biological replicates of each time point were stained with one of the 4 unique hash antibodies. Cells were sorted using a FACSAria Fusion or FACSAria II equipped with a 100 μm nozzle (BD Biosciences).

### Single-cell RNA library preparation and sequencing

HSPC single-cell RNA-seq was performed using the 10X Genomics platform. The Chromium Next GEM single cell 3‘ reagent kits v3.1 were implemented to prepare the libraries, following the official instruction manual (https://www.10xgenomics.com/support/single-cell-gene-expression/documentation/steps/library-prep/chromium-single-cell-3-reagent-kits-user-guide-v-3-1-chemistry). Briefly, 10,000 Lin^-^ cKit^+^ cells were sorted and enriched for HSCs by sorting additional 3000-4000 Lin_-_ cKit^+^ CD150^+^ CD48^-^ CD34^-^ cells. Cells were super-loaded according to the manufacturer’s instructions up until the cDNA amplification step. 1 ul/sample of HTO primers was spiked into the cDNA amplification PCR, and cDNA was amplified according to the 10x Single Cell 3’ v3.1 protocol aiming for a targeted cell recovery of 500-6000 cells. Following PCR, cDNA cleanup was performed by using SPRI to separate the HTO-derived cDNAs (in the supernatant) from the mRNA-derived cDNAs (retained on beads). The cDNA fraction was processed according to the manufacturer’s protocol to generate the transcriptome library. The quality of the obtained cDNA library upon adapter ligation and sample index PCR was assessed on an Agilent Bioanalyzer High sensitivity chip. Library sequencing was performed on the Novaseq 6000 Illumina sequencing platform.

### Filtering longitudinal single-cell RNAseq dataset

The cellranger pipeline (version 3.1.0) was used to align all reads to the mm10 genome and count the coverage of each gene in each cell. Based on the hashtag barcodes, cells were assigned to their corresponding time point (control, 3h, 24h, or 72h) and batch (four batches per time point). Cells with multiple barcodes (multiplets) or missing barcodes (negatives) were removed from the dataset. In the resulting count matrix (cells x genes), cells with a high amount of mitochondrial genes (> 5%) or a low amount of unique genes (<700) were filtered out. After the filtering steps, the following number of cells was present in each of the respective time points: control - 2474, 3h - 1661, 24h - 3462, 72h - 2449.

### Clustering and cell type annotation

The 500 most highly variable genes (HVGs) were identified in the control subset using analytic Pearson residuals (Lause, Berens, and Kobak 2021). The control subset was subsetted for the 500 HVGs, and the counts were L2 normalized. Next, a neighborhood graph was computed using 10 out of 50 principal components and the 15 nearest neighbors. The Leiden algorithm (resolution = 0.8) identified 14 distinct clusters in the control subset (Traag, Waltman, and van Eck 2019). Each cluster was appointed to a cell type based on 1) differentially expressed genes (DEGs) between the cluster of interest and all other clusters, 2) the expression profiles of the HVGs, 3) known marker genes and 4) correlation with cell types in a previously published dataset of the HSPCs (Nestorowa et al. 2016).

### Label transfer and UMAP representation

We identified the top 2000 HVGs in each subset and subsetted the complete dataset with the combined list of HVGs. Afterward, the dataset was L2 normalized, and the different subsets were integrated using Scanorama (Hie, Bryson, and Berger 2019). All 100 Scanorama-reduced dimensions were used to calculate a neighborhood graph (nearest neighbors = 15). A twodimensional UMAP representation was computed using the neighborhood graph. To transfer the cell type labels from the control subset to the response subsets (3h, 24h, and 72h), cells in the response subsets would adopt the cell type label that was most common among their 15 nearest neighbors (Euclidean distance) in the control subset. The integrated data was only used for label transfer and visualization purposes. For other downstream analyses, we use the filtered-only dataset. In this dataset, we removed the eosinophils and monocytes because of the small number of cells assigned to those cell types (10 and 52 respectively).

### Calculating gene set scores

The filtered dataset was L2 normalized and scaled to unit variance, and zero mean. The ISG score was calculated by subtracting the average expression of a random set of reference genes from the average expression of about 400 known ISGs (Scanpy function score_genes). Similarly, the stemness (Giladi et al. 2018), necroptosis (GO:0070266), pyroptosis (GO:0070269), myeloid TF (Kwok et al. 2020), monocyte and neutrophil differentiation, cell cycle (Giladi et al. 2018) and purine nucleotide synthesis (Vogel et al. 2019) score were calculated. Genes for each signature are available as a.csv file (Supplementary Table 1). Necroptosis and pyroptosis gene sets were retrieved from the Mouse Genome Database (MGD), Mouse Genome Informatics, The Jackson Laboratory, Bar Harbor, Maine. World Wide Web (URL: http://www.informatics.jax.org). (The data was retrieved in the year 2022)

### Differential abundance analysis

We used the R package Milo to perform an abundance analysis on the L2 normalized, filtered dataset (Dann et al. 2022). A neighborhood graph was built using 30 out of 100 of the Scanorama-reduced dimensions (see *Label transfer and UMAP representation*) and 30 nearest neighbors. Afterward, we followed the steps described in the accompanying tutorial (*Milo example on mouse gastrulation dataset*) for each response subset (3h, 24h, and 72h). In each analysis, the control subset served as the reference, to which the response subset would be compared.

### Identifying response genes

We used the edgeR-LRT method in the Libra R package to find the differentially expressed genes (DEGs) between the control and any of the response subsets, in each cluster (Robinson et al., 2010; Squair et al., 2021). We considered only DEGs with an adjusted p-value higher than 0.05 and a log-fold change higher than 1 in at least one cluster. For the downstream analyses of the response genes, we consider only the 500 DEGs with the highest p-values. In case a DEG is found in more than one cluster and/or time point, we take only the highest p-value into consideration.

### Change score

We L2 normalized the filtered dataset per cell. For each response gene (*i*) we took the mean expression (*μ*) in each cluster (*j*) per time point (*t*). The expression change was calculated as the absolutes sum of the derivative of the mean expression across all time points (here *m =* 3 for control-3h, 3h-24h, 24h-72h). Thus the change score (*c_i,j_*) per cluster for each response gene: (1)

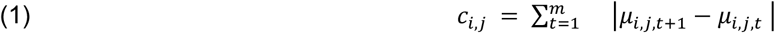

The result is a matrix with change scores per cluster for each of the response genes. We applied hierarchical clustering and grouped the response genes into 14 groups by setting a threshold at the cophenetic distance of 3 (Scipy function `cluster.hierarchy.linkage` and `cluster.hierarchy.fcluster`).

### Similarity score

We define the similarity score between the cluster-specific expression profiles of gene *i* and the expression profile of the same gene in the complete dataset in two steps. First, both the expression of the cluster and the complete dataset were scaled between 0 and 1 by min-max normalization (with *n* = 4 indicating the number of time points in the time series).

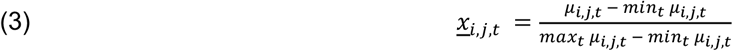

Note that *max_t_*, *μ_i,j,t_* and *min_t_ μ_i,j,t_* indicate the max/min of the mean expression for gene *i* cluster *j* among all time points *t*. Second, we calculate a dissimilarity score between the cluster and the whole data set for each response gene by subtracting the cluster-specific expression changes from the expression changes in the complete dataset. Third, the magnitude of these differences were summed and normalized by the number of time points. Finally, we turn the dissimilarity into a similarity score (*s_i,j_*) by subtracting it from 1 such that 1 presents complete similarity to the average behavior of all cells in the dataset.

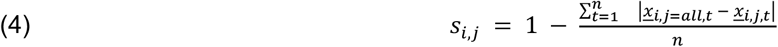

The result is a matrix with similarity scores per cluster for each of the response genes.

### Pseudotemporal ordering of cells during response

We opt to find a pseudotime axis that correlates with the actual arrow of time. Thus, we look for a transformation (*W* of size [*G*, 1]) of the expression data from all time points that reconstructs the experimental time point of each cell with minimal error (*ε*):

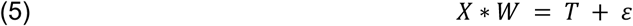

Here, *X* (of size [*N*, *G*]) is the filtered count matrix after L2 normalization, scaling and subsetting for all response genes. *T* is a vector (of size [*G*, 1]) with an (experimental) time assignment for each cell, created by taking the experimental time points (control, 3h, 24h, and 72h) and converting those to 0, 1, 2, and 3 respectively (alternatively one could consider using the actual time values on a log-scale). The least squares solution for *W* (which minimizes *ε^T^* * *ε*) is given by:

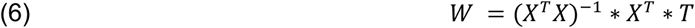

We used the expression matrix *X* with the size of 9983 cells and 2501 genes and the cells’ corresponding time labels to solve the above linear regression problem. We note that in order to avoid over-parametrization and to ensure the identifiability of the solution, the number of cells has to be larger than the number of genes. After *W* has been retrieved, a pseudotime coordinate can be calculated for each cell by:

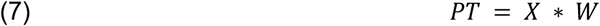

Where *PT* (of size [*N*, 1]) is the vector with the pseudotime coordinate for each cell. After pseudotime ordering of the cells, we smoothed the expression of candidate genes (e.g. the response genes) by taking the mean expression per 100 cells over the pseudotime axis. The expression values were scaled between 0 and 1 by min-max normalization. The 500 response genes were then clustered into 16 pseudotemporal expression patterns, using hierarchical clustering (threshold at cophenetic distance of 5.2).

### Gene expression in pseudotime

Expression profiles of individual genes in response pseudotime were derived using a combination of bin smoothing and bootstrapping. To find the expression profile in the complete dataset, bin smoothing with a 600-cell window size was performed on a sample of 50% of the cells in the dataset. This was repeated 20 times to find the mean expression, which defines the expression profile. The 95% confidence intervals were calculated by multiplying the standard error with 1.96 and subtracting or adding to the mean. For the cluster-specific expression profiles of individual genes a 50-cell window size was chosen instead, because of the smaller number of cells in each cluster.

### Gene score in pseudotime

The gene set score profile in response pseudotime was calculated using a combination of locally weighted least squares regression (LOESS) smoothing and bootstrapping. For each cluster LOESS smoothing with a first order regression model was applied to 50% of the cells. This was repeated 30 times. The score profile was derived by taking the mean and 1.96 times the standard error for the 95% confidence intervals.

## Code availability

All scripts used in this study are available on Github: https://github.com/bibouman/pri_HSPC.

## Data availability

The single-cell RNA-seq data were deposited in the Gene Expression Omnibus (GEO) under accession code GSE226824.

## Acknowledgement

The authors thank the staff at the DKFZ Flow Cytometry Core Facility, the DKFZ single cell Open Lab, and the DKFZ Animal Laboratory Services for their assistance and expertise. The authors thank Michael Milsom from the DKFZ for providing *Bax*^-/-^*Bak*^-/-^ mice. This work was supported by research funding from the Dietmar Hopp Foundation, and SFB873 funded by the Deutsche Forschungsgemeinschaft (DFG) (M.A.G.E.). B.J.B. and L.H. are supported by the Max-Delbrück-Center for Molecular Medicine in the Helmholtz Association (MDC), Berlin Institute for Medical Systems Biology (BIMSB). L.H. is also supported by the Bundesministerium für Bildung und Forschung (BMBF) “LeukoSyStem” consortium grant 01ZX1911B.

## Author information

Contributions: B.J.B, Y.D., L.H., and M.A.G.E. conceived and designed the study; Y.D., S.S., and M.A.G.E. designed the single-cell experimental setup; Y.D., S.S., F.P., A.R.I. and A.K. acquired the data; B.J.B. and L.H. developed the analysis pipeline; B.J.B., Y.D., L.H., and M.A.G.E. analyzed and interpreted data; F.G. and S.H. assisted with the analysis; B.J.B., Y.D., L.H., and M.A.G.E. drafted the manuscript; and all authors revised the manuscript for important intellectual content and approved the final version submitted for publication.

## Ethics declarations

Competing interests: The authors declare no competing financial interest.

