## Supplementary information for "Single-cell time series analysis reveals the dynamics of *in vivo* HSPC responses to inflammation"

### Supplementary Figures

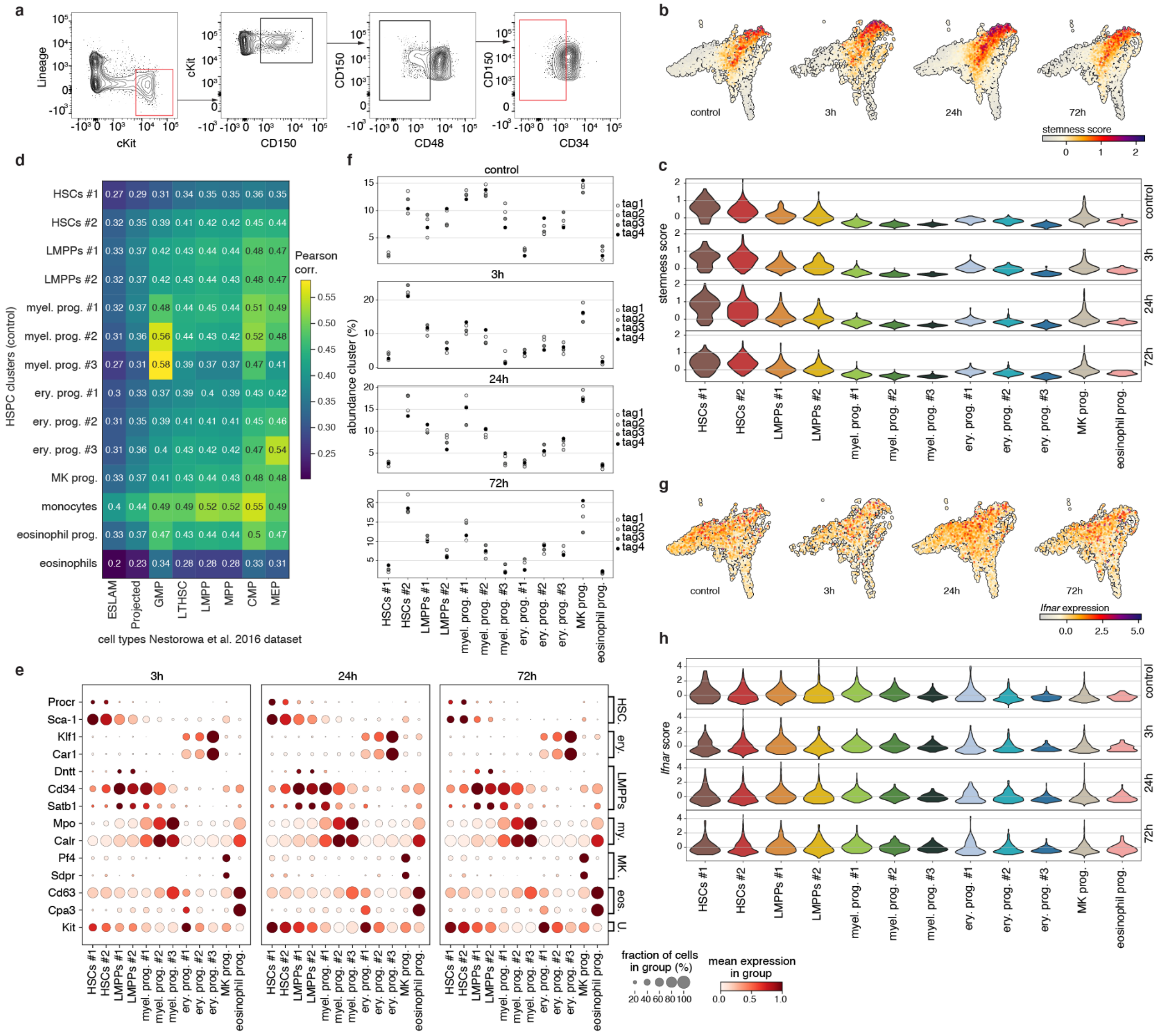


**Extended Fig. 1 | Characterization of cell clusters in the single-cell HSPC dataset. a**, Gating strategy for the single-cell RNA sequencing experiment, where 10,000 Lin$^-$ cKit$^+$ cells were sorted and enriched with 3000-5000 HSCs (Lin$^-$ Sca-1$^+$ cKit$^+$ CD150$^+$ CD48$^-$ CD34$^-$). Cells sorted are within the red gates **b,c,** UMAP projection (**b**) and violin plot (**c**) of stemness score (see Methods) in the IFNα-treated or control (PBS) subsets **d**, Pearson correlation between gene expression of clusters in our HSPC dataset versus the cell types in the HSPC dataset from *Nestorowa et al.* 2016. **e,** Expression of marker genes in the different clusters in the 3h, 24h, and 72h IFNα-treated subsets. erythroid (ery.); myeloid (my.); eosinophils (eos.); Universal (U.). **f**, Relative abundances of each cluster for the 4 different hashtags (biological replicates) in each timepoint. **g,h,** UMAP projection (**g**), and violin plot (**h**) of *Ifnar* expression in the four subsets.

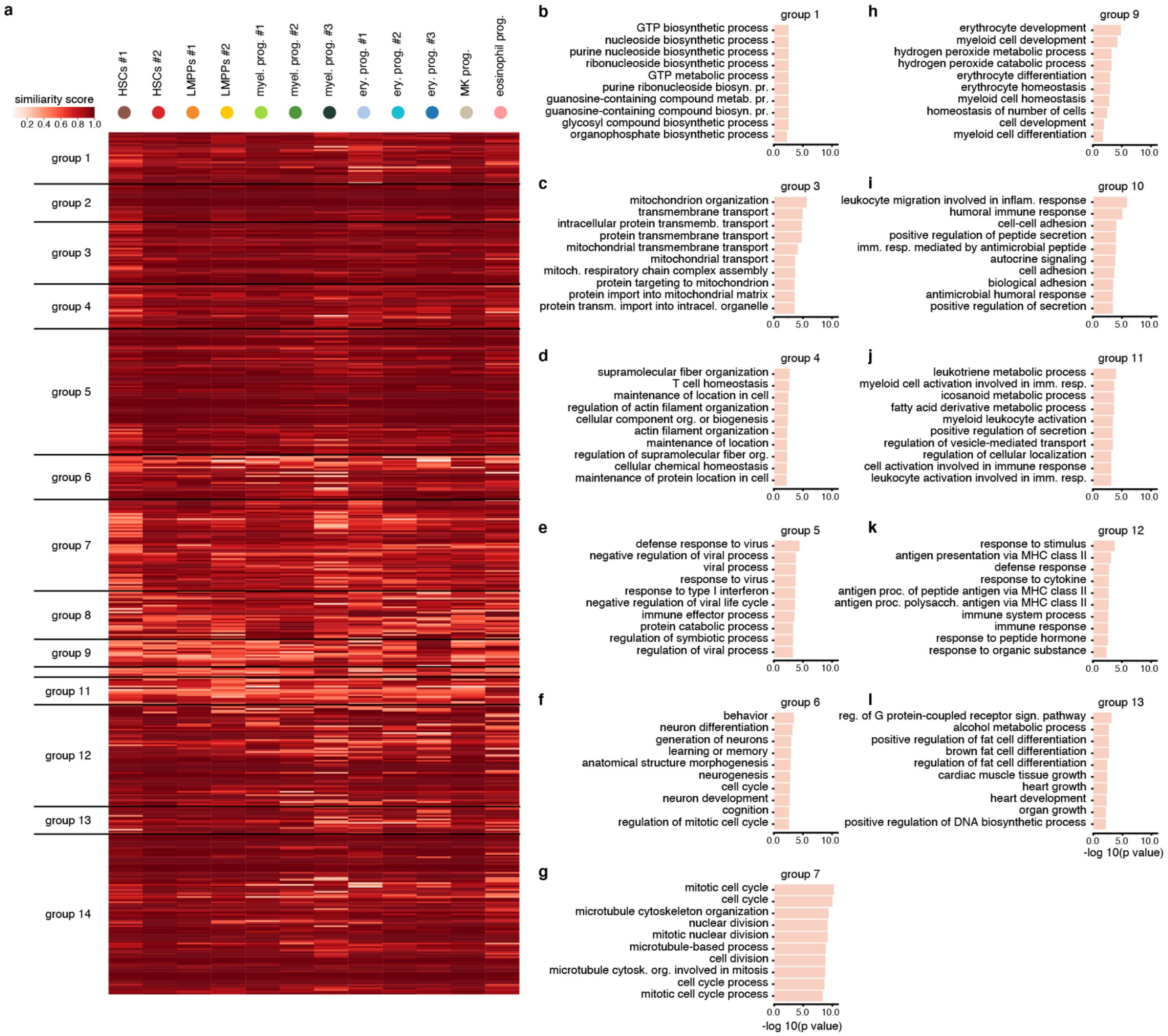


**Extended Fig. 2 | Inter-cluster similarity score analysis confirms the validity of cluster-specific inflammation signatures. a,** Similarity score (between 0 and 1) illustrating the similarity between the pattern of a gene in a specific cluster and the average pattern in the whole dataset. Genes are grouped as in Fig. 3b. **b-l,** GO terms significantly enriched in the different change score groups (as defined in Fig. 3b). The length of each bar represents the statistical significance of each term.

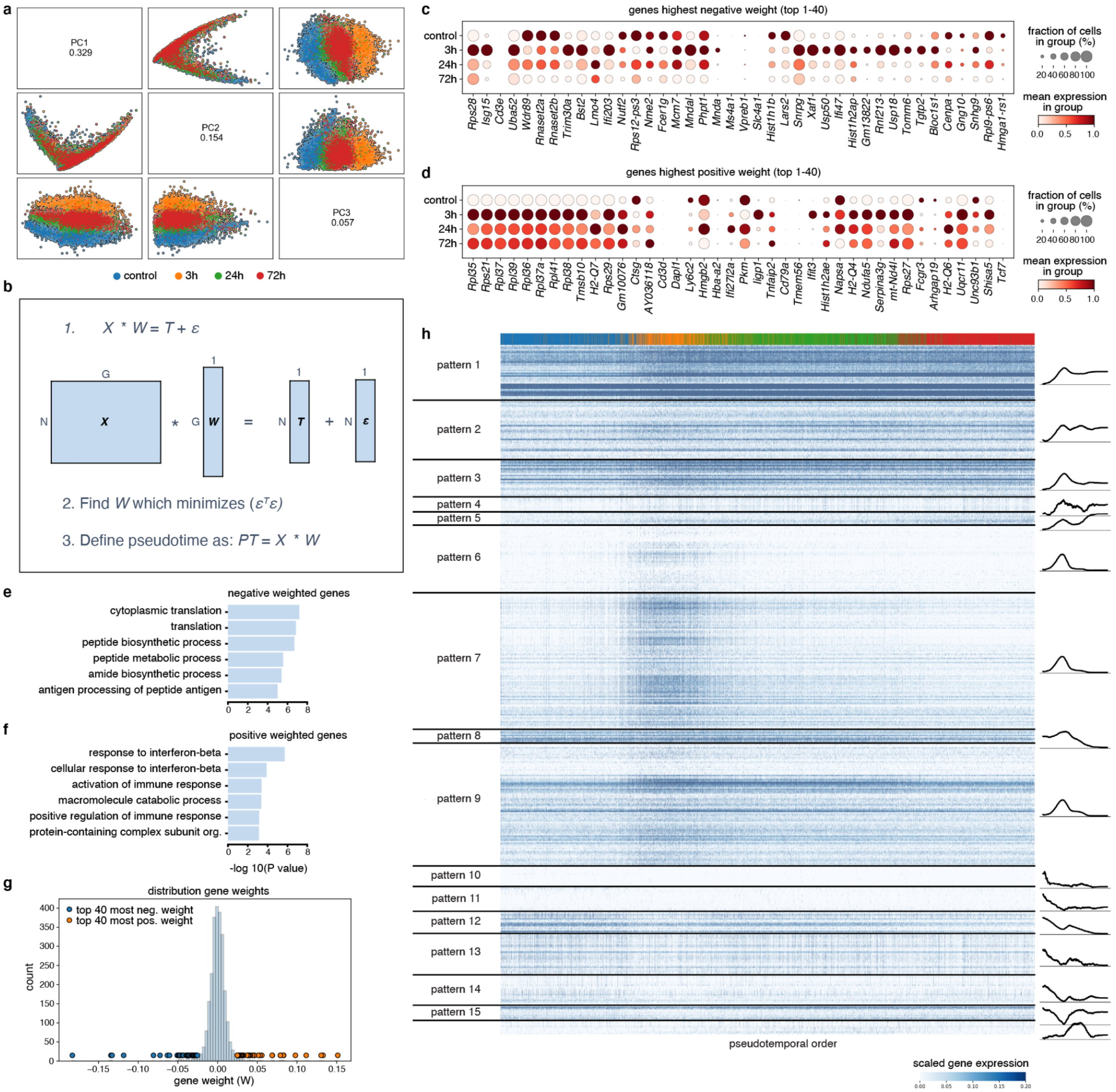


**Extended Fig. 3 | The response-pseudotime ordering of cells.** **a,** Principal components 1-3 of the non-batch corrected HSPC dataset. **b,** The response-pseudotime for each cell is calculated by solving a linear regression for the weight matrix W that transforms the data matrix X (with N cells and G genes) to the experimental time labels vector T, with minimal error (ε). **c,d,** gene expression of the top 40 genes with the lowest negative association (i.e., weight in the W matrix) with pseudotime (**c**), gene expression of the top 40 genes with the highest positive association with pseudotime (**d**) **e,f,** GO terms associated with negative weighted genes (**e**) and positive weighted genes (**f**). The length of each bar represents the statistical significance of each term. **g**, all weights in vector W, represented as histogram with points representing the top 40 most positive weighted genes (orange) and top 40 most negative weighted genes (blue). **h,** Non-smoothed expression of the top 500 DEGs. Expression patterns were grouped as in Fig. 4c.

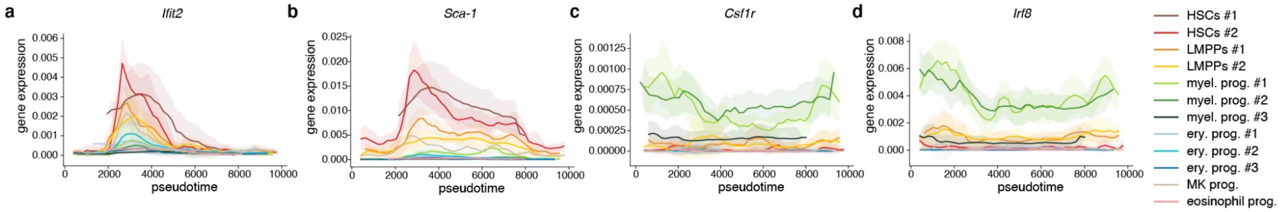


**Extended Fig. 4 | A response pseudotime analysis reveals gene dynamics in HSPCs following IFNα treatment.** **a-d,** Pseudotemporal expression of HSC-specific genes *Ifit2* (**a**) and *Sca-1* (**b**) and myeloid-specific genes *Csf1r* (**c**) and *Irf8* (**d**) in the different clusters.

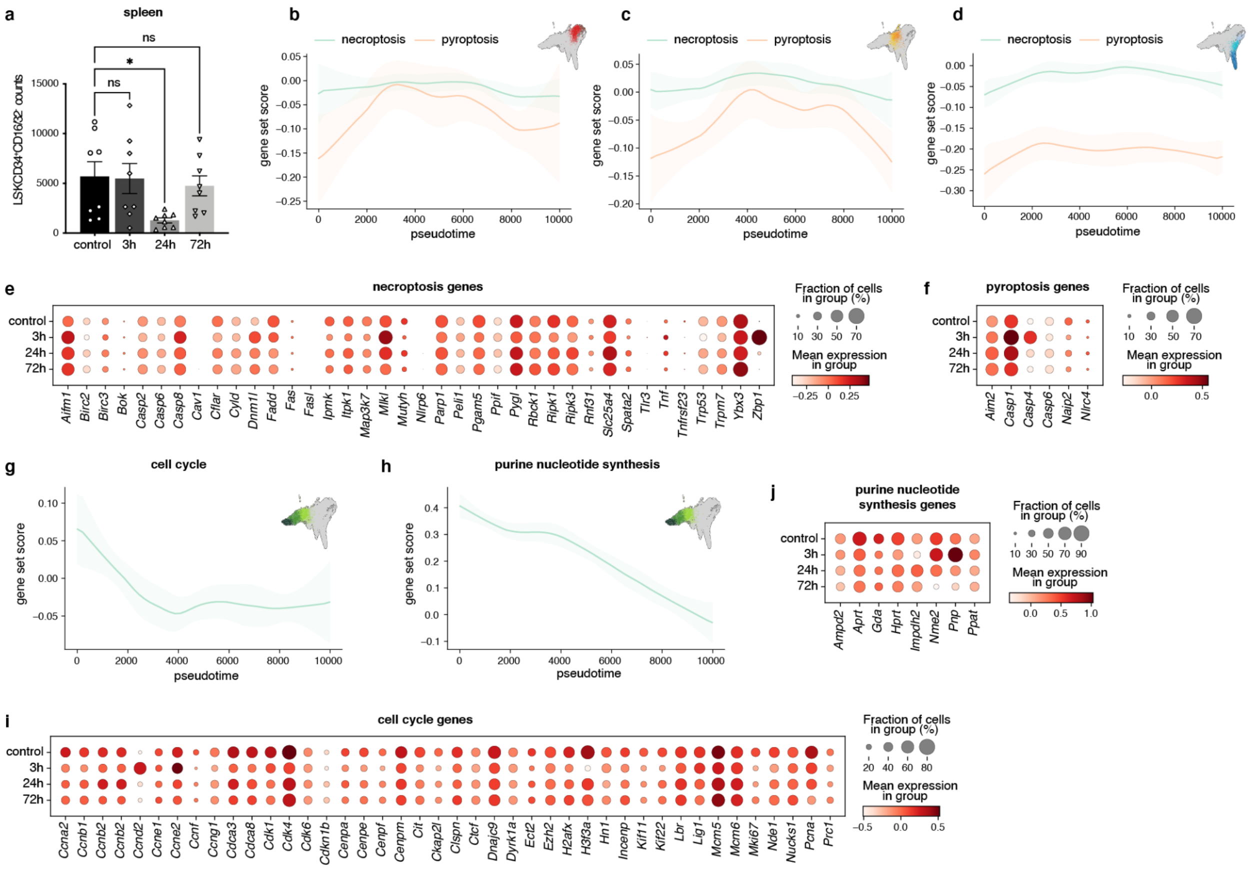

**Extended Fig. 5 | Reduction of myeloid progenitors functional properties following IFNα treatment. a,** Flow cytometric analysis of frequency of myeloid progenitors (Lin$^-$ Sca-1$^-$ cKit$^+$ CD34$^+$ CD16/32$^+$) in WT mice at 3h, 24h and 72h following IFNα or control (PBS) treatment in spleen. n= 8 biological replicates. **b-d,** Score of necroptosis gene signature and pyroptosis gene signature in all HSPCs (**b**), LMPPs (**c**) and erythroid progenitors (**d**) plotted in pseudotime. **e, f,** Expression of necroptosis **(e)** and pyroptosis **(f)** associated genes in myeloid progenitors in the four experimental subsets. **g,h,** Score of cell cycle **(g)** and purine nucleotide synthesis **(h)** genes in all three myeloid progenitor clusters plotted in pseudotime. **i, j,** Expression of cell cycle **(i)** and purine nucleotide synthesis **(j)** associated genes in myeloid progenitors in the four experimental subsets. Statistical significance in **a** was determined by an ordinary one-way ANOVA using Holm-Šídák's multiple comparisons test and at least two independent experiments were performed for a, and b only performed once; *P≤0.05, ***P≤ 0.001 ****P<0.0001.

Supplementary Note for:

### "Single-cell time series analysis reveals the dynamics of *in vivo* HSPC responses to inflammation"

Brigitte Joanne Bouman[*], Yasmin Demerdash[*], Shubhankar Sood, Florian Grünschläger, Franziska Pilz, Abdul Rahman Itani, Andrea Kuck, Simon Haas, Laleh Haghverdi[†], Marieke Alida Gertruda Essers[†]

* These authors contributed equally to this work.

† To whom correspondence should be addressed.

#### Recovering RNA velocity in single-cell time series

RNA velocity has emerged as a powerful tool to infer the transcriptional dynamics of single cells by leveraging the splicing information contained in the RNA sequencing data. The principle of RNA velocity is based on the fact that newly transcribed RNA molecules contain unspliced introns that are removed during the splicing process, whereas pre-existing RNA molecules lack these introns. By comparing the relative abundance of spliced and unspliced RNA molecules within each cell, RNA velocity can estimate the direction and speed of the transcriptional changes, which reflect the cell's developmental trajectory and/or fate. However, when applying RNA velocity on multiple datasets (such as a time series), several challenges arise. One of the main challenges is to choose the same gene set across all datasets, which is essential for making a fair comparison. Due to the unreliable quantification of unspliced versus spliced mRNA counts (i.e. input data for velocity estimation), many of the genes in each dataset will not pass quality controls and will therefore not be included in the estimation of RNA velocity. However, if the goal is to compare the velocities between datasets, it is important that the velocity vectors are made up from the same gene set components. One approach is to select the genes that pass quality control in all datasets, but this could lead to the omission of genes that are only activated in specific datasets and thus would not have sufficient unspliced counts in the other datasets. Additionally, there could be genes that pass quality control in one dataset but are filtered out in another dataset due to differences in data quality. Even if a gene passes quality control in all datasets, the inferred dynamics could still be different due to variations in data quality and insufficient robustness of the parameter fitting procedure, that can result in very different dynamical curves fit to very similar phase portrait data from the same gene, across different time points (see Figure 1). Therefore, choosing the same gene set for multiple datasets is not straightforward and requires careful consideration. These challenges highlight the need for more robust RNA velocity algorithms that can account for dataset-specific variations and enable fair comparisons between different datasets. Additionally, datasets with higher sequencing depth and better coverage full length RNAs could help overcome some of the issues related to quality control.

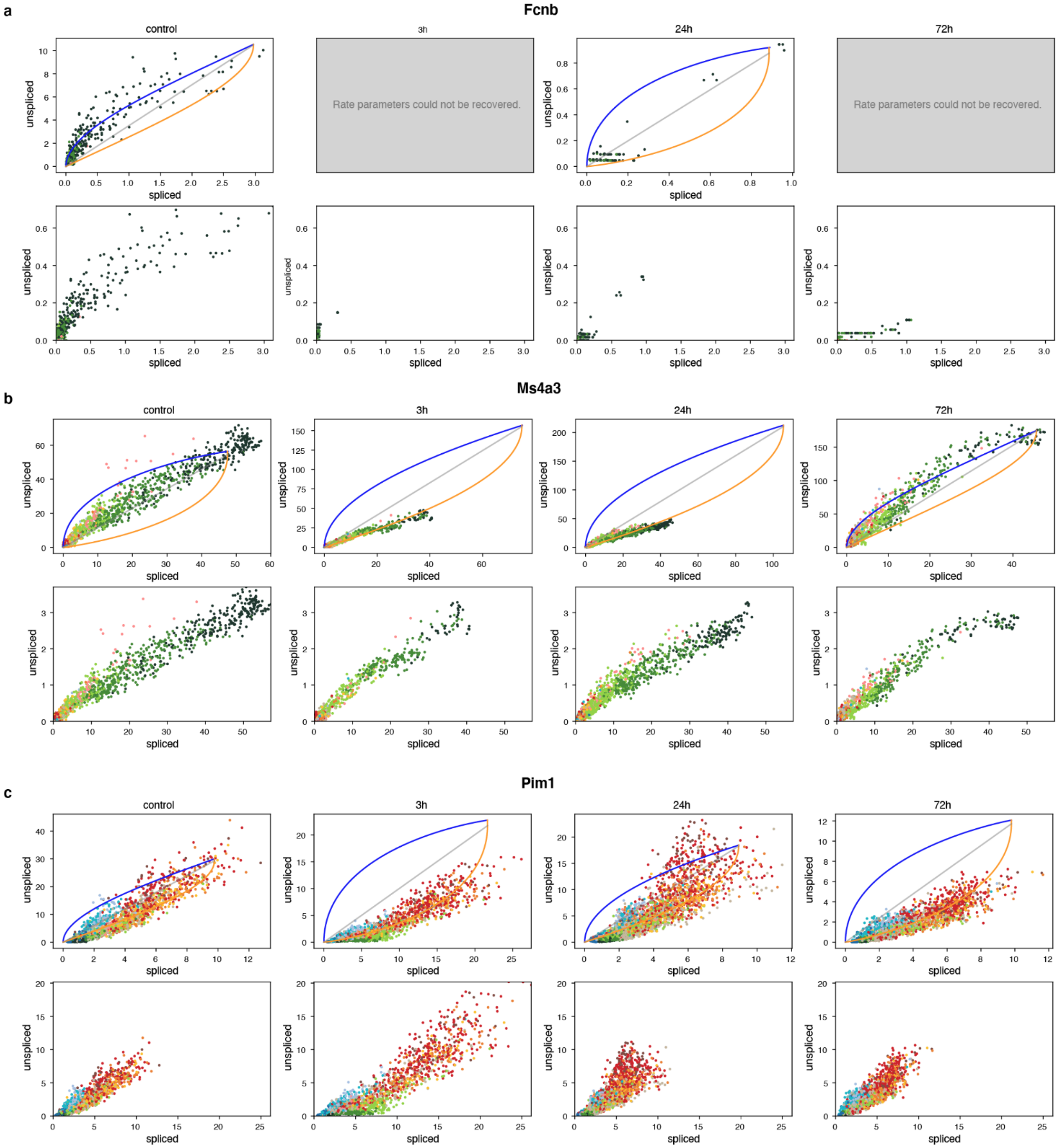


**Figure 1 |** Phase portrait of inferred dynamics of gene Fcnb (**a**), *Ms4a3* (**b**) and Pim1 (**c**) for the four different subsets in the HSPC dataset (control and three response subsets) (top row) and the imputed counts they were inferred from (bottom row). The figures show examples where the rate parameters could not be recovered (**a**), the rate parameters falsely predict a downregulation in some of the subsets (**b**), the counted unspliced and spliced counts are off for one of the subsets (3h) (**c**).

#### Supplementary Table 1

| Extended File | |
|---|---|
| **Gene** | Deescription |
| **Abca9** | Interferon stimulated gene (ISG) |
| **Abce1** | Interferon stimulated gene (ISG) |
| **Ablim3** | Interferon stimulated gene (ISG) |
| **Abtb2** | Interferon stimulated gene (ISG) |
| **Acsl1** | Interferon stimulated gene (ISG) |
| **Adamdec1** | Interferon stimulated gene (ISG) |
| **Adar** | Interferon stimulated gene (ISG) |
| **Adm** | Interferon stimulated gene (ISG) |
| **Agpat9** | Interferon stimulated gene (ISG) |
| **Aim2** | Interferon stimulated gene (ISG) |
| **Akt3** | Interferon stimulated gene (ISG) |
| **Aldh1a1** | Interferon stimulated gene (ISG) |
| **Alyref** | Interferon stimulated gene (ISG) |
| **Amph** | Interferon stimulated gene (ISG) |
| **Angptl1** | Interferon stimulated gene (ISG) |
| **Ankrd22** | Interferon stimulated gene (ISG) |
| **Apol2** | Interferon stimulated gene (ISG) |
| **Apol6** | Interferon stimulated gene (ISG) |
| **Aqp9** | Interferon stimulated gene (ISG) |
| **Arg2** | Interferon stimulated gene (ISG) |
| **Arhgef3** | Interferon stimulated gene (ISG) |
| **Arntl** | Interferon stimulated gene (ISG) |
| **Atf2** | Interferon stimulated gene (ISG) |
| **Atf3** | Interferon stimulated gene (ISG) |

| | |
|---|---|
| **B2m** | Interferon stimulated gene (ISG) |
| **Bag1** | Interferon stimulated gene (ISG) |
| **Bak1** | Interferon stimulated gene (ISG) |
| **Banf1** | Interferon stimulated gene (ISG) |
| **Batf2** | Interferon stimulated gene (ISG) |
| **Bax** | Interferon stimulated gene (ISG) |
| **Bcl2** | Interferon stimulated gene (ISG) |
| **Bcl2l1** | Interferon stimulated gene (ISG) |
| **Bcl3** | Interferon stimulated gene (ISG) |
| **Birc2** | Interferon stimulated gene (ISG) |
| **Birc3** | Interferon stimulated gene (ISG) |
| **Blvra** | Interferon stimulated gene (ISG) |
| **Blzf1** | Interferon stimulated gene (ISG) |
| **Bst2** | Interferon stimulated gene (ISG) |
| **Bub1** | Interferon stimulated gene (ISG) |
| **C10orf10** | Interferon stimulated gene (ISG) |
| **C15orf48** | Interferon stimulated gene (ISG) |
| **C1S** | Interferon stimulated gene (ISG) |
| **C22orf28** | Interferon stimulated gene (ISG) |
| **C4orf32** | Interferon stimulated gene (ISG) |
| **C4orf33** | Interferon stimulated gene (ISG) |
| **C9orf91** | Interferon stimulated gene (ISG) |
| **Calr** | Interferon stimulated gene (ISG) |
| **Canx** | Interferon stimulated gene (ISG) |
| **Casp1** | Interferon stimulated gene (ISG) |
| **Casp7** | Interferon stimulated gene (ISG) |

| | |
|---|---|
| **Ccdc75** | Interferon stimulated gene (ISG) |
| **Ccl11** | Interferon stimulated gene (ISG) |
| **Ccl2** | Interferon stimulated gene (ISG) |
| **Ccl22** | Interferon stimulated gene (ISG) |
| **Ccl4** | Interferon stimulated gene (ISG) |
| **Ccl5** | Interferon stimulated gene (ISG) |
| **Ccna1** | Interferon stimulated gene (ISG) |
| **Ccr1** | Interferon stimulated gene (ISG) |
| **Ccr7** | Interferon stimulated gene (ISG) |
| **Cd163** | Interferon stimulated gene (ISG) |
| **Cd274** | Interferon stimulated gene (ISG) |
| **Cd38** | Interferon stimulated gene (ISG) |
| **Cd40** | Interferon stimulated gene (ISG) |
| **Cd69** | Interferon stimulated gene (ISG) |
| **Cd74** | Interferon stimulated gene (ISG) |
| **Cd9** | Interferon stimulated gene (ISG) |
| **Cdk17** | Interferon stimulated gene (ISG) |
| **Cdk18** | Interferon stimulated gene (ISG) |
| **Cdkn1a** | Interferon stimulated gene (ISG) |
| **Ces1** | Interferon stimulated gene (ISG) |
| **Cfb** | Interferon stimulated gene (ISG) |
| **Chmp5** | Interferon stimulated gene (ISG) |
| **Chuk** | Interferon stimulated gene (ISG) |
| **Ciita** | Interferon stimulated gene (ISG) |
| **Clec4d** | Interferon stimulated gene (ISG) |
| **Clec4e** | Interferon stimulated gene (ISG) |

| | |
|---|---|
| **Clec5a** | Interferon stimulated gene (ISG) |
| **Cnp** | Interferon stimulated gene (ISG) |
| **Commd3** | Interferon stimulated gene (ISG) |
| **Cpt1a** | Interferon stimulated gene (ISG) |
| **Creb3l3** | Interferon stimulated gene (ISG) |
| **Crebbp** | Interferon stimulated gene (ISG) |
| **Crebzf** | Interferon stimulated gene (ISG) |
| **Crp** | Interferon stimulated gene (ISG) |
| **Cry1** | Interferon stimulated gene (ISG) |
| **Csrnp1** | Interferon stimulated gene (ISG) |
| **Cx3cl1** | Interferon stimulated gene (ISG) |
| **Cxcl10** | Interferon stimulated gene (ISG) |
| **Cxcl9** | Interferon stimulated gene (ISG) |
| **Cxcr4** | Interferon stimulated gene (ISG) |
| **Cyp1b1** | Interferon stimulated gene (ISG) |
| **Cyth1** | Interferon stimulated gene (ISG) |
| **Dcp1a** | Interferon stimulated gene (ISG) |
| **Ddit4** | Interferon stimulated gene (ISG) |
| **Ddx58** | Interferon stimulated gene (ISG) |
| **Ddx60** | Interferon stimulated gene (ISG) |
| **Dhx58** | Interferon stimulated gene (ISG) |
| **Dtx3l** | Interferon stimulated gene (ISG) |
| **Duox2** | Interferon stimulated gene (ISG) |
| **Dusp5** | Interferon stimulated gene (ISG) |
| **Dynlt1** | Interferon stimulated gene (ISG) |
| **Ehd4** | Interferon stimulated gene (ISG) |

| | |
|---|---|
| **Eif2ak2** | Interferon stimulated gene (ISG) |
| **Eif3l** | Interferon stimulated gene (ISG) |
| **Elf1** | Interferon stimulated gene (ISG) |
| **Enpp1** | Interferon stimulated gene (ISG) |
| **Epas1** | Interferon stimulated gene (ISG) |
| **Erlin1** | Interferon stimulated gene (ISG) |
| **Etv6** | Interferon stimulated gene (ISG) |
| **Ext1** | Interferon stimulated gene (ISG) |
| **Fadd** | Interferon stimulated gene (ISG) |
| **Fam125b** | Interferon stimulated gene (ISG) |
| **Fam134b** | Interferon stimulated gene (ISG) |
| **Fam46a** | Interferon stimulated gene (ISG) |
| **Fam46c** | Interferon stimulated gene (ISG) |
| **Fam70a** | Interferon stimulated gene (ISG) |
| **Fbxo6** | Interferon stimulated gene (ISG) |
| **Fcgr1a** | Interferon stimulated gene (ISG) |
| **Ffar2** | Interferon stimulated gene (ISG) |
| **Fgr** | Interferon stimulated gene (ISG) |
| **Fkbp5** | Interferon stimulated gene (ISG) |
| **Flt1** | Interferon stimulated gene (ISG) |
| **Fndc3b** | Interferon stimulated gene (ISG) |
| **Fndc4** | Interferon stimulated gene (ISG) |
| **Fosl1** | Interferon stimulated gene (ISG) |
| **Fut4** | Interferon stimulated gene (ISG) |
| **Fv1** | Interferon stimulated gene (ISG) |
| **Fzd5** | Interferon stimulated gene (ISG) |

| | |
|---|---|
| **G6Pc** | Interferon stimulated gene (ISG) |
| **Gak** | Interferon stimulated gene (ISG) |
| **Galnt2** | Interferon stimulated gene (ISG) |
| **Gbp2** | Interferon stimulated gene (ISG) |
| **Gbp4** | Interferon stimulated gene (ISG) |
| **Gbp5** | Interferon stimulated gene (ISG) |
| **Gca** | Interferon stimulated gene (ISG) |
| **Gch1** | Interferon stimulated gene (ISG) |
| **Gem** | Interferon stimulated gene (ISG) |
| **Gja4** | Interferon stimulated gene (ISG) |
| **Gk** | Interferon stimulated gene (ISG) |
| **Glipr2** | Interferon stimulated gene (ISG) |
| **Glrx** | Interferon stimulated gene (ISG) |
| **Gmpr** | Interferon stimulated gene (ISG) |
| **Gpx2** | Interferon stimulated gene (ISG) |
| **Gtpbp2** | Interferon stimulated gene (ISG) |
| **Gzmb** | Interferon stimulated gene (ISG) |
| **Hbxip** | Interferon stimulated gene (ISG) |
| **Heg1** | Interferon stimulated gene (ISG) |
| **Herc6** | Interferon stimulated gene (ISG) |
| **Hesx1** | Interferon stimulated gene (ISG) |
| **Hk2** | Interferon stimulated gene (ISG) |
| **Hla-f** | Interferon stimulated gene (ISG) |
| **Hla-g** | Interferon stimulated gene (ISG) |
| **Hnrnpul1** | Interferon stimulated gene (ISG) |
| **Hpse** | Interferon stimulated gene (ISG) |

| | |
|---|---|
| **Hsh2d** | Interferon stimulated gene (ISG) |
| **Hyal1** | Interferon stimulated gene (ISG) |
| **Hyal2** | Interferon stimulated gene (ISG) |
| **Hyal3** | Interferon stimulated gene (ISG) |
| **Ido1** | Interferon stimulated gene (ISG) |
| **Ifi27l2** | Interferon stimulated gene (ISG) |
| **Ifi30** | Interferon stimulated gene (ISG) |
| **Ifi35** | Interferon stimulated gene (ISG) |
| **Ifi44** | Interferon stimulated gene (ISG) |
| **Ifi44l** | Interferon stimulated gene (ISG) |
| **Ifih1** | Interferon stimulated gene (ISG) |
| **Ifit1** | Interferon stimulated gene (ISG) |
| **Ifit2** | Interferon stimulated gene (ISG) |
| **Ifit3** | Interferon stimulated gene (ISG) |
| **Ifitm1** | Interferon stimulated gene (ISG) |
| **Ifitm2** | Interferon stimulated gene (ISG) |
| **Ifitm3** | Interferon stimulated gene (ISG) |
| **Ifne** | Interferon stimulated gene (ISG) |
| **Igfbp2** | Interferon stimulated gene (ISG) |
| **Ikbkb** | Interferon stimulated gene (ISG) |
| **Ikbke** | Interferon stimulated gene (ISG) |
| **Ikbkg** | Interferon stimulated gene (ISG) |
| **Il10** | Interferon stimulated gene (ISG) |
| **Il12B** | Interferon stimulated gene (ISG) |
| **Il12rb1** | Interferon stimulated gene (ISG) |
| **Il15** | Interferon stimulated gene (ISG) |

| | |
|---|---|
| **Il15ra** | Interferon stimulated gene (ISG) |
| **Il17rb** | Interferon stimulated gene (ISG) |
| **Il1r1** | Interferon stimulated gene (ISG) |
| **Il1rn** | Interferon stimulated gene (ISG) |
| **Il23a** | Interferon stimulated gene (ISG) |
| **Il23r** | Interferon stimulated gene (ISG) |
| **Il28ra** | Interferon stimulated gene (ISG) |
| **Il6** | Interferon stimulated gene (ISG) |
| **Il6st** | Interferon stimulated gene (ISG) |
| **Impa2** | Interferon stimulated gene (ISG) |
| **Irf1** | Interferon stimulated gene (ISG) |
| **Irf2** | Interferon stimulated gene (ISG) |
| **Irf3** | Interferon stimulated gene (ISG) |
| **Irf7** | Interferon stimulated gene (ISG) |
| **Irf9** | Interferon stimulated gene (ISG) |
| **Isg15** | Interferon stimulated gene (ISG) |
| **Isg20** | Interferon stimulated gene (ISG) |
| **Itch** | Interferon stimulated gene (ISG) |
| **Ivns1abp** | Interferon stimulated gene (ISG) |
| **Jak2** | Interferon stimulated gene (ISG) |
| **Jun** | Interferon stimulated gene (ISG) |
| **Junb** | Interferon stimulated gene (ISG) |
| **Lamp3** | Interferon stimulated gene (ISG) |
| **Lap3** | Interferon stimulated gene (ISG) |
| **Lcn2** | Interferon stimulated gene (ISG) |
| **Lepr** | Interferon stimulated gene (ISG) |

| | |
|---|---|
| **Lgals3** | Interferon stimulated gene (ISG) |
| **Lgals9** | Interferon stimulated gene (ISG) |
| **Lgmn** | Interferon stimulated gene (ISG) |
| **Lipa** | Interferon stimulated gene (ISG) |
| **Lmo2** | Interferon stimulated gene (ISG) |
| **Lta** | Interferon stimulated gene (ISG) |
| **Ly6e** | Interferon stimulated gene (ISG) |
| **Mab21l2** | Interferon stimulated gene (ISG) |
| **Mafb** | Interferon stimulated gene (ISG) |
| **Maff** | Interferon stimulated gene (ISG) |
| **Map3k14** | Interferon stimulated gene (ISG) |
| **Map3k5** | Interferon stimulated gene (ISG) |
| **Mapkapk2** | Interferon stimulated gene (ISG) |
| **Mastl** | Interferon stimulated gene (ISG) |
| **Mavs** | Interferon stimulated gene (ISG) |
| **Max** | Interferon stimulated gene (ISG) |
| **Mb21d1** | Interferon stimulated gene (ISG) |
| **Mcl1** | Interferon stimulated gene (ISG) |
| **Med14** | Interferon stimulated gene (ISG) |
| **Mfn1** | Interferon stimulated gene (ISG) |
| **Micb** | Interferon stimulated gene (ISG) |
| **Mkx** | Interferon stimulated gene (ISG) |
| **Mov10** | Interferon stimulated gene (ISG) |
| **Ms4a4a** | Interferon stimulated gene (ISG) |
| **Mst1r** | Interferon stimulated gene (ISG) |
| **Mt1h** | Interferon stimulated gene (ISG) |

| | |
|---|---|
| **Mthfd2l** | Interferon stimulated gene (ISG) |
| **Myd88** | Interferon stimulated gene (ISG) |
| **Myof** | Interferon stimulated gene (ISG) |
| **N4bp1** | Interferon stimulated gene (ISG) |
| **Nampt** | Interferon stimulated gene (ISG) |
| **Napa** | Interferon stimulated gene (ISG) |
| **Ncf1** | Interferon stimulated gene (ISG) |
| **Ncoa3** | Interferon stimulated gene (ISG) |
| **Ndc80** | Interferon stimulated gene (ISG) |
| **Nfil3** | Interferon stimulated gene (ISG) |
| **Nlrx1** | Interferon stimulated gene (ISG) |
| **Nmi** | Interferon stimulated gene (ISG) |
| **Nod2** | Interferon stimulated gene (ISG) |
| **Nos2** | Interferon stimulated gene (ISG) |
| **Npas2** | Interferon stimulated gene (ISG) |
| **Nt5c3** | Interferon stimulated gene (ISG) |
| **Nup50** | Interferon stimulated gene (ISG) |
| **Oas1** | Interferon stimulated gene (ISG) |
| **Oas1b** | Interferon stimulated gene (ISG) |
| **Oas2** | Interferon stimulated gene (ISG) |
| **Oas3** | Interferon stimulated gene (ISG) |
| **Oasl** | Interferon stimulated gene (ISG) |
| **Odc1** | Interferon stimulated gene (ISG) |
| **Ogfr** | Interferon stimulated gene (ISG) |
| **Optn** | Interferon stimulated gene (ISG) |
| **Otub1** | Interferon stimulated gene (ISG) |

| | |
|---|---|
| **Otub2** | Interferon stimulated gene (ISG) |
| **P2ry6** | Interferon stimulated gene (ISG) |
| **Pabpc4** | Interferon stimulated gene (ISG) |
| **Padi2** | Interferon stimulated gene (ISG) |
| **Pdgfrl** | Interferon stimulated gene (ISG) |
| **Pdia3** | Interferon stimulated gene (ISG) |
| **Pdk1** | Interferon stimulated gene (ISG) |
| **Pfdn6** | Interferon stimulated gene (ISG) |
| **Pfkfb3** | Interferon stimulated gene (ISG) |
| **Phf15** | Interferon stimulated gene (ISG) |
| **Pi4k2b** | Interferon stimulated gene (ISG) |
| **Pias1** | Interferon stimulated gene (ISG) |
| **Pim3** | Interferon stimulated gene (ISG) |
| **Pin1** | Interferon stimulated gene (ISG) |
| **Plekha4** | Interferon stimulated gene (ISG) |
| **Plin2** | Interferon stimulated gene (ISG) |
| **Plp1** | Interferon stimulated gene (ISG) |
| **Plscr1** | Interferon stimulated gene (ISG) |
| **Pml** | Interferon stimulated gene (ISG) |
| **Pmm2** | Interferon stimulated gene (ISG) |
| **Pnpt1** | Interferon stimulated gene (ISG) |
| **Pnrc1** | Interferon stimulated gene (ISG) |
| **Ppm1k** | Interferon stimulated gene (ISG) |
| **Prkd2** | Interferon stimulated gene (ISG) |
| **Prkra** | Interferon stimulated gene (ISG) |
| **Psmb5** | Interferon stimulated gene (ISG) |

| | |
|---|---|
| **Psmb6** | Interferon stimulated gene (ISG) |
| **Psmb8** | Interferon stimulated gene (ISG) |
| **Psmb9** | Interferon stimulated gene (ISG) |
| **Ptpn2** | Interferon stimulated gene (ISG) |
| **Ptpn6** | Interferon stimulated gene (ISG) |
| **Pus1** | Interferon stimulated gene (ISG) |
| **Pxk** | Interferon stimulated gene (ISG) |
| **Rab27a** | Interferon stimulated gene (ISG) |
| **Raf1** | Interferon stimulated gene (ISG) |
| **Rasgef1b** | Interferon stimulated gene (ISG) |
| **Rassf4** | Interferon stimulated gene (ISG) |
| **Rbck1** | Interferon stimulated gene (ISG) |
| **Rbm25** | Interferon stimulated gene (ISG) |
| **Rela** | Interferon stimulated gene (ISG) |
| **Rgs1** | Interferon stimulated gene (ISG) |
| **Ripk1** | Interferon stimulated gene (ISG) |
| **Rnase4** | Interferon stimulated gene (ISG) |
| **Rnasel** | Interferon stimulated gene (ISG) |
| **Rnf114** | Interferon stimulated gene (ISG) |
| **Rnf216** | Interferon stimulated gene (ISG) |
| **Rpl22** | Interferon stimulated gene (ISG) |
| **Rps15a** | Interferon stimulated gene (ISG) |
| **Rsad2** | Interferon stimulated gene (ISG) |
| **Rtp4** | Interferon stimulated gene (ISG) |
| **S100a8** | Interferon stimulated gene (ISG) |
| **Saa1** | Interferon stimulated gene (ISG) |

| | |
|---|---|
| **Samd4a** | Interferon stimulated gene (ISG) |
| **Samhd1** | Interferon stimulated gene (ISG) |
| **Sat1** | Interferon stimulated gene (ISG) |
| **Scarb2** | Interferon stimulated gene (ISG) |
| **Sco2** | Interferon stimulated gene (ISG) |
| **Sectm1** | Interferon stimulated gene (ISG) |
| **Serpinb9** | Interferon stimulated gene (ISG) |
| **Serpine1** | Interferon stimulated gene (ISG) |
| **Serping1** | Interferon stimulated gene (ISG) |
| **Sike1** | Interferon stimulated gene (ISG) |
| **Sirpa** | Interferon stimulated gene (ISG) |
| **Slc15a3** | Interferon stimulated gene (ISG) |
| **Slc16a1** | Interferon stimulated gene (ISG) |
| **Slc1a1** | Interferon stimulated gene (ISG) |
| **Slc25a28** | Interferon stimulated gene (ISG) |
| **Slc25a30** | Interferon stimulated gene (ISG) |
| **Slfn5** | Interferon stimulated gene (ISG) |
| **Smad3** | Interferon stimulated gene (ISG) |
| **Snn** | Interferon stimulated gene (ISG) |
| **Socs1** | Interferon stimulated gene (ISG) |
| **Socs2** | Interferon stimulated gene (ISG) |
| **Socs3** | Interferon stimulated gene (ISG) |
| **Sp110** | Interferon stimulated gene (ISG) |
| **Spaca3** | Interferon stimulated gene (ISG) |
| **Spn** | Interferon stimulated gene (ISG) |
| **Spsb1** | Interferon stimulated gene (ISG) |

| | |
|---|---|
| **Sptlc2** | Interferon stimulated gene (ISG) |
| **Ssbp3** | Interferon stimulated gene (ISG) |
| **Stap1** | Interferon stimulated gene (ISG) |
| **Stard5** | Interferon stimulated gene (ISG) |
| **Stat1** | Interferon stimulated gene (ISG) |
| **Stat2** | Interferon stimulated gene (ISG) |
| **Stat3** | Interferon stimulated gene (ISG) |
| **Steap4** | Interferon stimulated gene (ISG) |
| **Sun2** | Interferon stimulated gene (ISG) |
| **Tagap** | Interferon stimulated gene (ISG) |
| **Tank** | Interferon stimulated gene (ISG) |
| **Tap1** | Interferon stimulated gene (ISG) |
| **Tap2** | Interferon stimulated gene (ISG) |
| **Tapbp** | Interferon stimulated gene (ISG) |
| **Tbk1** | Interferon stimulated gene (ISG) |
| **Tbx3** | Interferon stimulated gene (ISG) |
| **Tcf7l2** | Interferon stimulated gene (ISG) |
| **Tdrd7** | Interferon stimulated gene (ISG) |
| **Tfec** | Interferon stimulated gene (ISG) |
| **Thbd** | Interferon stimulated gene (ISG) |
| **Ticam1** | Interferon stimulated gene (ISG) |
| **Timp1** | Interferon stimulated gene (ISG) |
| **Tlk2** | Interferon stimulated gene (ISG) |
| **Tlr3** | Interferon stimulated gene (ISG) |
| **Tlr7** | Interferon stimulated gene (ISG) |
| **Tlr8** | Interferon stimulated gene (ISG) |

| | |
|---|---|
| **Tmem140** | Interferon stimulated gene (ISG) |
| **Tmem173** | Interferon stimulated gene (ISG) |
| **Tmem51** | Interferon stimulated gene (ISG) |
| **Tnf** | Interferon stimulated gene (ISG) |
| **Tnfaip3** | Interferon stimulated gene (ISG) |
| **Tnfrsf10a** | Interferon stimulated gene (ISG) |
| **Tnfrsf9** | Interferon stimulated gene (ISG) |
| **Tnfsf10** | Interferon stimulated gene (ISG) |
| **Traf2** | Interferon stimulated gene (ISG) |
| **Traf3** | Interferon stimulated gene (ISG) |
| **Traf6** | Interferon stimulated gene (ISG) |
| **Trafd1** | Interferon stimulated gene (ISG) |
| **Trex1** | Interferon stimulated gene (ISG) |
| **Trim14** | Interferon stimulated gene (ISG) |
| **Trim21** | Interferon stimulated gene (ISG) |
| **Trim25** | Interferon stimulated gene (ISG) |
| **Trim38** | Interferon stimulated gene (ISG) |
| **Trim5** | Interferon stimulated gene (ISG) |
| **Trim56** | Interferon stimulated gene (ISG) |
| **Txnip** | Interferon stimulated gene (ISG) |
| **Tyk2** | Interferon stimulated gene (ISG) |
| **Tymp** | Interferon stimulated gene (ISG) |
| **Uba7** | Interferon stimulated gene (ISG) |
| **Ube2l6** | Interferon stimulated gene (ISG) |
| **Ulk4** | Interferon stimulated gene (ISG) |
| **Unc93b1** | Interferon stimulated gene (ISG) |

| | |
|---|---|
| **Upp2** | Interferon stimulated gene (ISG) |
| **Uri1** | Interferon stimulated gene (ISG) |
| **Usp18** | Interferon stimulated gene (ISG) |
| **Vamp5** | Interferon stimulated gene (ISG) |
| **Vav1** | Interferon stimulated gene (ISG) |
| **Vegfc** | Interferon stimulated gene (ISG) |
| **Vmp1** | Interferon stimulated gene (ISG) |
| **Wars** | Interferon stimulated gene (ISG) |
| **Whamm** | Interferon stimulated gene (ISG) |
| **Xaf1** | Interferon stimulated gene (ISG) |
| **Xcl1** | Interferon stimulated gene (ISG) |
| **Xpr1** | Interferon stimulated gene (ISG) |
| **Zbp1** | Interferon stimulated gene (ISG) |
| **Zc3hav1** | Interferon stimulated gene (ISG) |
| **Znf295** | Interferon stimulated gene (ISG) |
| **Znf385b** | Interferon stimulated gene (ISG) |
| **1110017F19Rik** | Stemness gene |
| **2410006H16Rik;Snord65** | Stemness gene |
| **2900062L11Rik;6530401D17Rik** | Stemness gene |
| **4931406C07Rik** | Stemness gene |
| **AK046388** | Stemness gene |
| **AK079675** | Stemness gene |
| **AK197603;Uba7;AK157941;Cdh29;Ube1l;D330022A01Rik;Cdhr4;Ip6k1** | Stemness gene |
| **Acot1** | Stemness gene |
| **Aldoc** | Stemness gene |
| **B144;Lst1** | Stemness gene |
| **Basp1** | Stemness gene |
| **Bgn** | Stemness gene |
| **Car2** | Stemness gene |
| **Cd27** | Stemness gene |
| **Cd274** | Stemness gene |

| | |
|---|---|
| **Cd74** | Stemness gene |
| **Cd81** | Stemness gene |
| **Cish** | Stemness gene |
| **Clip3** | Stemness gene |
| **Dapp1** | Stemness gene |
| **Dkkl1** | Stemness gene |
| **Eltd1** | Stemness gene |
| **Fchsd2;mKIAA0769** | Stemness gene |
| **Gabarapl1** | Stemness gene |
| **Gcnt2** | Stemness gene |
| **Gimap1** | Stemness gene |
| **Gimap6** | Stemness gene |
| **Gm6251** | Stemness gene |
| **Gpr56** | Stemness gene |
| **H2-K1** | Stemness gene |
| **H2-T10;H2-T22;H2-T9;H2-t9;H2-T23** | Stemness gene |
| **Hlf** | Stemness gene |
| **Hoxb2** | Stemness gene |
| **Ifitm1** | Stemness gene |
| **Ifitm2** | Stemness gene |
| **Ifitm3** | Stemness gene |
| **Krt18** | Stemness gene |
| **Ldhb** | Stemness gene |
| **Leprel2** | Stemness gene |
| **Lhcgr** | Stemness gene |
| **Lmo2** | Stemness gene |
| **Ly6a** | Stemness gene |
| **Malat1** | Stemness gene |
| **Mecom** | Stemness gene |
| **Mllt3** | Stemness gene |
| **Mpl** | Stemness gene |
| **Msi2;msi2;LOC100504473** | Stemness gene |
| **Mycn** | Stemness gene |
| **Myct1** | Stemness gene |
| **Myl10** | Stemness gene |
| **Nfe2** | Stemness gene |
| **Oasl2** | Stemness gene |
| **Osbpl1a** | Stemness gene |
| **Pcp4l1** | Stemness gene |
| **Pde4b** | Stemness gene |

| | |
|---|---|
| **Pdzk1ip1** | Stemness gene |
| **Pglyrp2;tagL** | Stemness gene |
| **Pla2g16** | Stemness gene |
| **Pnrc1** | Stemness gene |
| **Procr** | Stemness gene |
| **Ptplad2** | Stemness gene |
| **Ptpn18** | Stemness gene |
| **Ptprcap** | Stemness gene |
| **Rbp1** | Stemness gene |
| **Rbpms;Rbpms2** | Stemness gene |
| **Rgs1** | Stemness gene |
| **Rtp4** | Stemness gene |
| **Serpina3f;Serpina3g** | Stemness gene |
| **Shisa5** | Stemness gene |
| **Srgn** | Stemness gene |
| **Stxbp4** | Stemness gene |
| **Tbxas1** | Stemness gene |
| **Tgtp2;Gm12185** | Stemness gene |
| **Tmem176a** | Stemness gene |
| **Tmem176b** | Stemness gene |
| **Tnip3** | Stemness gene |
| **Txnip** | Stemness gene |
| **Ube2l6** | Stemness gene |
| **Wfdc2** | Stemness gene |
| **Whamm;mKIAA1971** | Stemness gene |
| **Zfand5** | Stemness gene |
| **Zfp608;mKIAA1281;Znf608;AK155149** | Stemness gene |
| **Zfp831** | Stemness gene |
| **Aifm1** | Necroptosis |
| **Birc2** | Necroptosis |
| **Birc3** | Necroptosis |
| **Bok** | Necroptosis |
| **Casp2** | Necroptosis |
| **Casp6** | Necroptosis |
| **Casp8** | Necroptosis |
| **Cav1** | Necroptosis |
| **Cflar** | Necroptosis |
| **Cyld** | Necroptosis |
| **Dnm1l** | Necroptosis |
| **Fadd** | Necroptosis |

| | |
|---|---|
| **Fas** | Necroptosis |
| **Fasl** | Necroptosis |
| **Ipmk** | Necroptosis |
| **Itpk1** | Necroptosis |
| **Map3k7** | Necroptosis |
| **Mlkl** | Necroptosis |
| **Mutyh** | Necroptosis |
| **Nlrp6** | Necroptosis |
| **Parp1** | Necroptosis |
| **Peli1** | Necroptosis |
| **Pgam5** | Necroptosis |
| **Ppif** | Necroptosis |
| **Pygl** | Necroptosis |
| **Rbck1** | Necroptosis |
| **Ripk1** | Necroptosis |
| **Ripk3** | Necroptosis |
| **Rnf31** | Necroptosis |
| **Slc25a4** | Necroptosis |
| **Spata2** | Necroptosis |
| **Tlr3** | Necroptosis |
| **Tnf** | Necroptosis |
| **Tnfrsf23** | Necroptosis |
| **Trp53** | Necroptosis |
| **Trpm7** | Necroptosis |
| **Ybx3** | Necroptosis |
| **Zbp1** | Necroptosis |
| **Aim2** | Pyroptosis |
| **Casp1** | Pyroptosis |
| **Casp4** | Pyroptosis |
| **Casp6** | Pyroptosis |
| **Dhx9** | Pyroptosis |
| **Nlrc4** | Pyroptosis |
| **Naip2** | Pyroptosis |
| **Papss2** | Monoccyte differenttiation |
| **Ass1** | Monoccyte differenttiation |
| **Tcfec** | Monoccyte differenttiation |
| **Trem2** | Monoccyte differenttiation |
| **Rassf4** | Monoccyte differenttiation |
| **Ly6c2** | Monoccyte differenttiation |
| **Ms4a6c** | Monoccyte differenttiation |

| | |
|---|---|
| **F13a1** | Monoccyte differenttiation |
| **Ctss** | Monoccyte differenttiation |
| **Klf4** | Monoccyte differenttiation |
| **S100a4** | Monoccyte differenttiation |
| **Slpi** | Monoccyte differenttiation |
| **Prdx4** | Monoccyte differenttiation |
| **Hpse** | Monoccyte differenttiation |
| **Tifab** | Monoccyte differenttiation |
| **Csf1r** | Monoccyte differenttiation |
| **Ly86** | Monoccyte differenttiation |
| **Emb** | Monoccyte differenttiation |
| **Glipr1** | Monoccyte differenttiation |
| **Irf8** | Monoccyte differenttiation |
| **Elane** | Monoccyte differenttiation |
| **Ifitm1** | Monoccyte differenttiation |
| **Gpr56** | Monoccyte differenttiation |
| **Cd34** | Monoccyte differenttiation |
| **Eltd1** | Monoccyte differenttiation |
| **Serpina3f** | Monoccyte differenttiation |
| **Ifitm6** | Neutrophil differentiation |
| **Chi3l3** | Neutrophil differentiation |
| **S100a9** | Neutrophil differentiation |
| **Ngp** | Neutrophil differentiation |
| **Syne1** | Neutrophil differentiation |
| **S100a8** | Neutrophil differentiation |
| **Orm1** | Neutrophil differentiation |
| **Chi3l1** | Neutrophil differentiation |
| **Ltf** | Neutrophil differentiation |
| **Lrg1** | Neutrophil differentiation |
| **Pglyrp1** | Neutrophil differentiation |
| **Itgb2l** | Neutrophil differentiation |
| **Camp** | Neutrophil differentiation |
| **Cd177** | Neutrophil differentiation |
| **Lcn2** | Neutrophil differentiation |
| **Fcnb** | Neutrophil differentiation |
| **Mpo** | Neutrophil differentiation |
| **Elane** | Neutrophil differentiation |
| **Gstm1** | Neutrophil differentiation |
| **Ifitm1** | Neutrophil differentiation |
| **Gpr56** | Neutrophil differentiation |

| | |
|---|---|
| **Cd34** | Neutrophil differentiation |
| **Eltd1** | Neutrophil differentiation |
| **Serpina3f** | Neutrophil differentiation |
| **Prc1** | cellcycle |
| **Pcna** | cellcycle |
| **Nucks1** | cellcycle |
| **Nde1** | cellcycle |
| **Mki67** | cellcycle |
| **Mcm6** | cellcycle |
| **Mcm5** | cellcycle |
| **Lig1** | cellcycle |
| **Lbr** | cellcycle |
| **Kif22** | cellcycle |
| **Kif11** | cellcycle |
| **Incenp** | cellcycle |
| **Hn1** | cellcycle |
| **H3f3a** | cellcycle |
| **H2afx** | cellcycle |
| **Ezh2** | cellcycle |
| **Ect2** | cellcycle |
| **Dyrk1a** | cellcycle |
| **Dnajc9** | cellcycle |
| **Ctcf** | cellcycle |
| **Clspn** | cellcycle |
| **Ckap2l** | cellcycle |
| **Cit** | cellcycle |
| **Cenpm** | cellcycle |
| **Cenpf** | cellcycle |
| **Cenpe** | cellcycle |
| **Cenpa** | cellcycle |
| **Cdk1** | cellcycle |
| **Cdca8** | cellcycle |
| **Cdca3** | cellcycle |
| **Ccnb2** | cellcycle |
| **Ccnd2** | cellcycle |
| **Ccne1** | cellcycle |
| **Ccne2** | cellcycle |
| **Cdkn1b** | cellcycle |
| **Ccnb1** | cellcycle |
| **Ccna2** | cellcycle |

| | |
|---|---|
| **Ccnf** | cellcycle |
| **Ccnb2** | cellcycle |
| **Ccng1** | cellcycle |
| **Cdk4** | cellcycle |
| **Cdk6'** | cellcycle |
| **Nme2** | Purine nuceleotide synthesis |
| **Pnp** | Purine nuceleotide synthesis |
| **Impdh2** | Purine nuceleotide synthesis |
| **Ppat** | Purine nuceleotide synthesis |
| **Hprt** | Purine nuceleotide synthesis |
| **Aprt** | Purine nuceleotide synthesis |
| **Gda** | Purine nuceleotide synthesis |
| **Ampd2** | Purine nuceleotide synthesis |
| **Cebpe** | Myeloid transcription factor |
| **Calr** | Myeloid transcription factor |
| **Arid3a** | Myeloid transcription factor |
| **Gfi1** | Myeloid transcription factor |
| **Lmo4** | Myeloid transcription factor |
| **Cebpa** | Myeloid transcription factor |
| **Spi1** | Myeloid transcription factor |
| **Cux1** | Myeloid transcription factor |
| **Scand1** | Myeloid transcription factor |
| **Nfkbia** | Myeloid transcription factor |
| **Irf8** | Myeloid transcription factor |
| **Id2** | Myeloid transcription factor |
| **Chd3** | Myeloid transcription factor |
| **Cbfa2t3** | Myeloid transcription factor |
| **Etv6** | Myeloid transcription factor |
| **Stat3** | Myeloid transcription factor |
| **Pnrc1** | Myeloid transcription factor |
| **Pbx1** | Myeloid transcription factor |
| **Mef2c** | Myeloid transcription factor |
| **Fli1** | Myeloid transcription factor |
| **Elf1** | Myeloid transcription factor |
| **Lmo2** | Myeloid transcription factor |
| **Cited2** | Myeloid transcription factor |
| **Sox4** | Myeloid transcription factor |
| **Runx1** | Myeloid transcription factor |
| **Gata2** | Myeloid transcription factor |
| **Nfe2** | Myeloid transcription factor |

| | |
|---|---|
| **Myb** | Myeloid transcription factor |
| **Foxp1** | Myeloid transcription factor |
| **Zfpm1** | Myeloid transcription factor |
| **Hmgb3** | Myeloid transcription factor |
| **Klf1** | Myeloid transcription factor |
| **Gfi1b** | Myeloid transcription factor |
| **Tcf3** | Myeloid transcription factor |
| **Pa2g4** | Myeloid transcription factor |
| **Mbd2** | Myeloid transcription factor |
| **Gata1** | Myeloid transcription factor |
| **Phf10** | Myeloid transcription factor |
| **Phb2** | Myeloid transcription factor |
| **Gtf2f1** | Myeloid transcription factor |
| **Csda** | Myeloid transcription factor |
| **E2f4** | Myeloid transcription factor |
| **Cited4** | Myeloid transcription factor |
| **Ccne1** | Myeloid transcription factor |
